# From comparative connectomics to large-scale working memory modeling in macaque and marmoset

**DOI:** 10.1101/2025.03.17.643781

**Authors:** Loïc Magrou, Panagiota Theodoni, Amy F. T. Arnsten, Marcello G. P. Rosa, Xiao-Jing Wang

**Author notes:** These authors contributed equally (Loïc Magrou, Panagiota Theodoni).

## Abstract

Although macaques and marmosets are both primates of choice for studying the brain mechanisms of cognition, they differ in key aspects of anatomy and behavior. Interestingly, recent connectomic analysis revealed that strong top-down projections from the prefrontal cortex to the posterior parietal cortex, present in macaques and important for executive function, are absent in marmosets. Here, we propose a consensus mapping that bridges the two species’ cortical atlases and allows for direct area-to-area comparison of their connectomes, which are then used to build comparative computational large-scale modeling of the frontoparietal circuit for working memory. We found that the macaque model exhibits resilience against distractors, a prerequisite for normal working memory function. By contrast, the marmoset model predicts a sensitivity to distractibility commonly observed behaviorally in this species. Surprisingly, this contrasting trend can be swapped by scaling intrafrontal and frontoparietal, and offer a credible prediction to the marmoset’s behavior in this specific task.

## INTRODUCTION

From Cuvier and Lamarck to Darwin and Wallace, comparative methods have historically been at the root of modern biology^1^. In neuroscience, recent advances of connectomics^2,3^, spanning from local^4–7^ to multiregional^8–16^ connectivity, offer a novel and quantitative approach to comparison between species. Comparative work on cortical connectivity has yielded important results in terms of graph theoretical^8,17,18^ and scaling properties of the brain^19–22^. However, whereas structural information is critical, it is often insufficient to predict dynamical behavior of a recurrent brain system, the understanding of which requires concomitant progress in physiological experiments and computational modeling^23^.

In this context, Working Memory (WM), the brain’s ability to internally maintain and manipulate information in the absence of sensory stimulation, has become a topic of focus, as its representation is distributed across multiple —-but not all –– brain regions^24–26^. In WM dependent tasks, neurons in certain parts of the macaque cortex show robust, self-sustained, information-specific, mnemonic persistent activity^27–30^. Experimental observations motivated connectome-based modeling of macaque monkey cortex for distributed WM^31,32^. Consistent with experimental observations^33^, the reciprocal loop between posterior parietal cortex (PPC) and prefrontal cortex (PFC), making together the frontoparietal network, was found to play a major role in WM. Furthermore, a salient characteristic of normal WM is its resilience against distractors^34–36^, whose electrophysiology is now well understood^37–39^, and which has been shown in modeling studies to critically depend on the PFC and its top-down influence onto the PPC in the macaque monkey cortex^31,32,40^.

Much less is known for the marmoset species, which is rapidly becoming another important animal model in our field^41–43^. Indeed, marmosets possess many of the cognitive attributes of both macaques and humans, whilst being at the same time faster to breed and easier to handle in experiments. Additionally, and non trivially for the question at hand here, they were shown to possess the same frontoparietal network known to be the substrate of WM in macaques and humans^44^. They are able to perform WM tasks^45^ and have been shown to exhibit sustained delay activity in the prefrontal cortex during those tasks^46^. Finally, the recent publication of the most complete retrograde tracer-based marmoset connectome available to date^8,9^ allows for analysis and comparison with the macaque.

The in-depth investigation of the marmoset’s WM capabilities is still at its early stage, but what seems to transpire from experimentalists is that, although they can perform such tasks, they are harder to train^47^, overall less capable to hold items in WM^48^ and remain significantly more distractible than macaques^49^. Through this convergence of anatomical, functional and behavioral evidence, we find ourselves with the unique opportunity to push forward the comparative method into the realm of computational neuroscience. We can now apply the same model to both species, knowing that any behavioral difference that might emerge will be solely due to the differences in anatomy. Specifically, differential distractibility could be captured with adequate modeling.

The present work tackles this important comparative question, with two main goals. We will first provide a common anatomical framework for the two species, which we have named “consensus mapping”, whose purpose will be to reduce the macaque and the marmoset independent atlases to a common parcelation scheme where areal connectivity can be compared directly, and then convert all the available anatomical data to it, connectivity and spine count gradient alike. Second, we will constrain an otherwise identical model with those two sets of now one-to-one comparable anatomical data and investigate for commonalities and differences in the model’s behavior.

## RESULTS

### Marmosets lack key fronto-parietal feedback from 46d 9/46d

The atlases of macaque and marmoset respectively hold 91 and 116 areas before consensus, with 40 and 55 injected areas (**Figure 1A and B**, injected areas in bold). Consensus brings the two atlases to 74 aras on both sides, capitalizing on primate analog broader areas. Applying consensus to connectivity itself, the number of injected areas are now 35 areas for the macaque and 45 for the marmoset. Of these, 29 are common to both species (**Figure 1C and D**, injected areas in bold).

**Figure 1.**
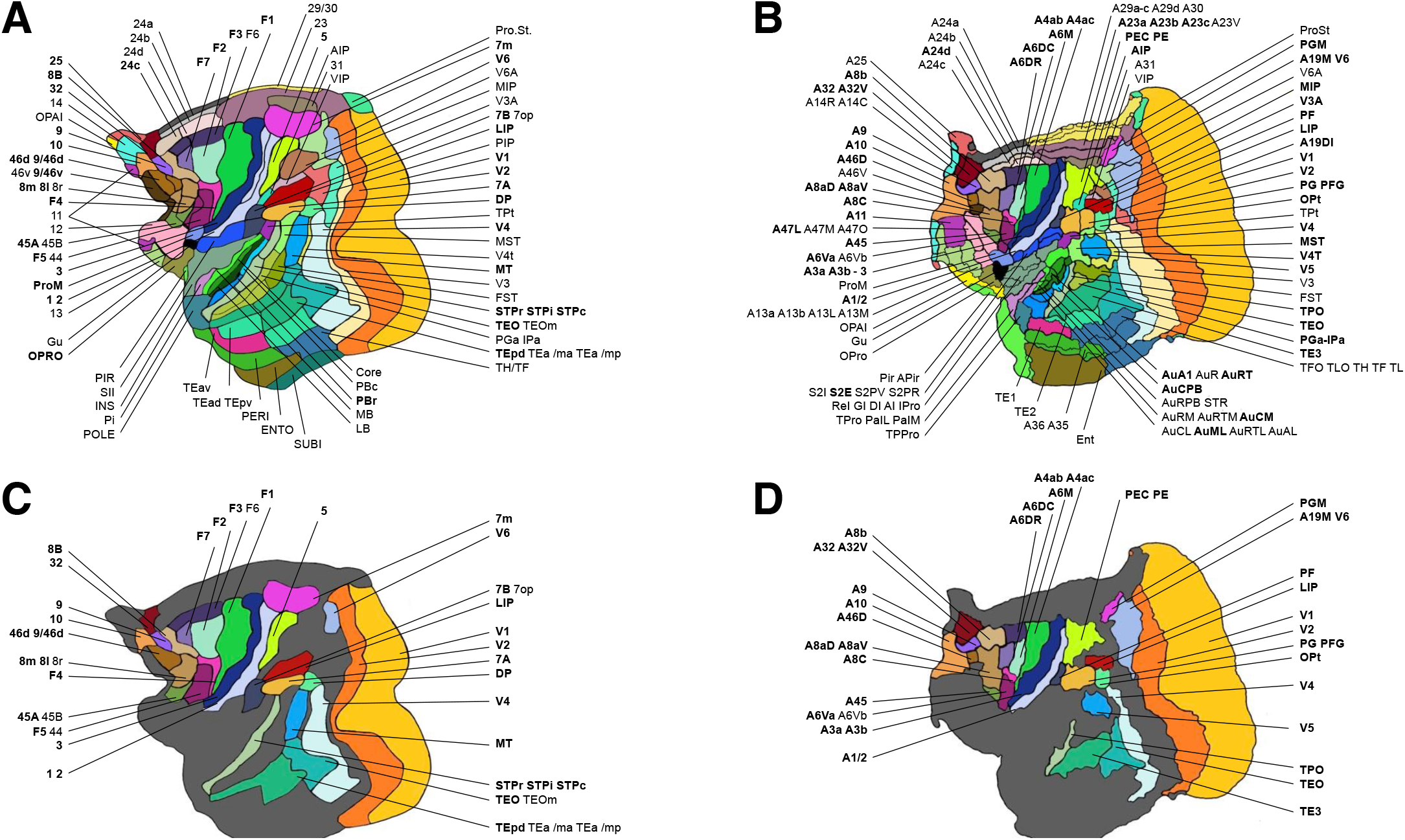
Flattened illustrations of the consensus mapping. (A) Macaque original 91 area atlas from Markov et al. ^14^. (B) Marmoset original 116 area atlas from Theodoni et al. ^8^. (C) Macaque 29 consensus areas commonly injected with the marmoset. (D) Marmoset 29 homolog consensus areas from (C), commonly injected with the macaque. Area names placed on the same line represent areas that will be aggregated as one area by the consensus mapping, and are shown as separate areas of identical color in (A) and (B), and then fused in (C) and (D). Names in bold are those for which there at least one injection the the original connectivity dataset. For easiness of read, area labels are identically ordered between macaque (A and C) and marmoset (B and D). The full table of consensus equivalence across the two species is available in **Table S3**.

The 29×29 square consensus connectivity matrices (**Figure 2**), comprise a total of 794 and 809 connections for the macaque and the marmoset respectively, over a total of 29 ×28 = 812 possible connections (self connections are ignored), leading to respective graph densities of 0.68 and 0.72. For the macaque, the distribution of log_10_ *FLN* (Fraction of Labeled Neurons, *i.e*. connection weights) is best approximated by a normal distribution, with a mean of − 2.60, which is slightly lower than that of the marmoset at − 2.28, which is consistent with previous results from Theodoni et al. ^8^ in spite of modifications produced by the consensus (**Figure 3A**). Also consistent with previous work, if the macaque spans the usual 6 orders of magnitude, as described elsewhere^14,50^, the marmoset spans only 5 of those orders^8,9^. Correlation between macaque and marmoset log_10_ *FLN* data is 0.59. Notably, The marmoset lacks some of the topdown connections from DLPFC to PPC that the macaque has, notably from area 46d 9/46d to LIP, which is a major contributor to spatial WM in the macaque. Conversely, the macaque appears to be missing some — albeit weak — connections from area 10 to medial parietal and posterior visual areas, suggesting that A10 in marmoset may retain some features of the more posterior prefrontal areas, such as the frontal eye field (A8aD A8aV) (**Figure 2B**).

**Figure 2.**
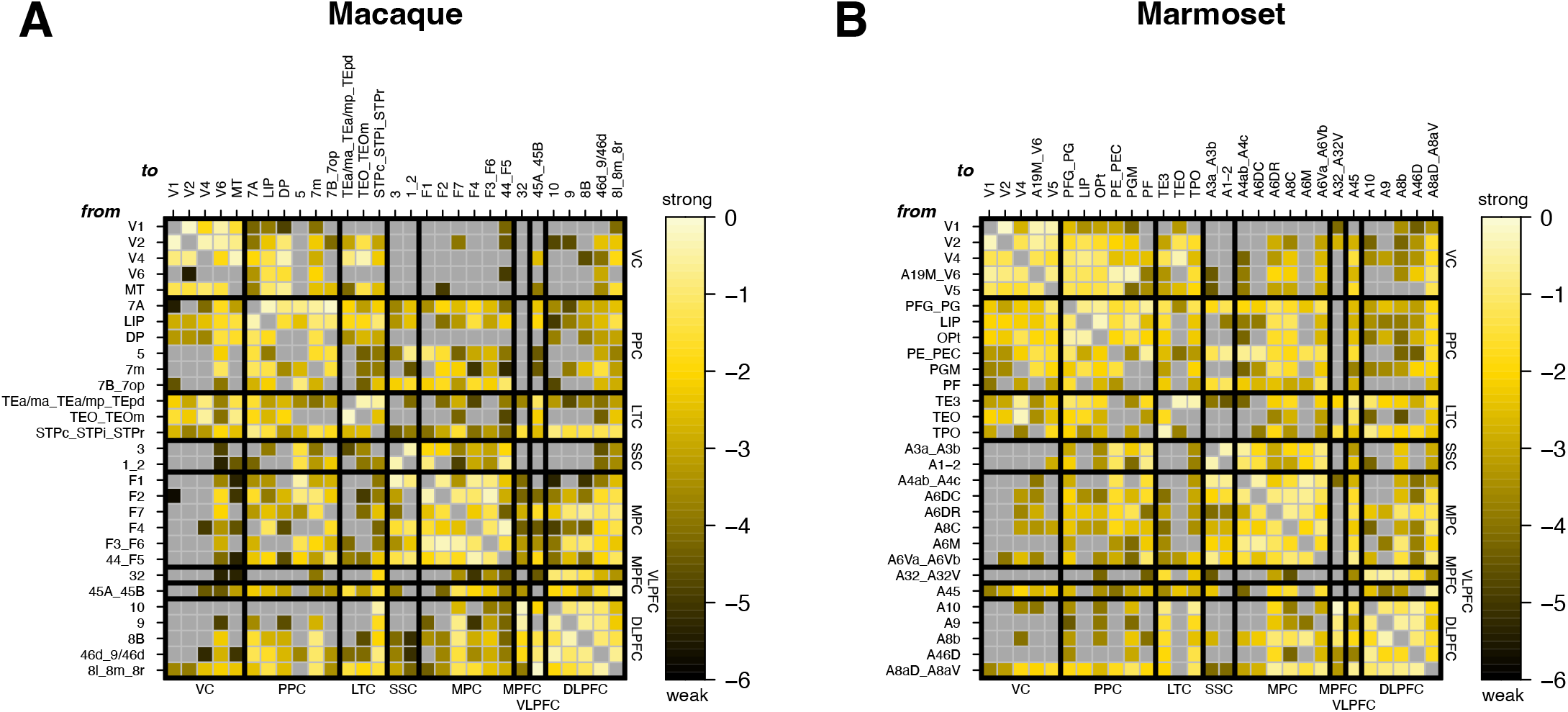
29×29 common consensus log_10_*FLN* matrices for both species. (A) Macaque (B) Marmoset. In these, each square of color represents a connection from source areas as rows to target areas as columns. Colors vary according to log_10_ *FLN* values, from very weak (black) to very strong (whitish yellow). Grey is the background color of the matrices and a grey square therefore identifies an absent connection. Black thicker lines separate brain regions, see **Table S3** for a key to regional acronyms. Areas merged by consensus are labeled by concatenating together the names of the original parcelation areas, separated by an underscore. Thanks to consensus, each connection are here one-to-one comparable across species. The corresponding SLN matrices are available in **Figure S4**, and a comprehensive connectivity table with both FLN and SLN values can be found in **Table S3**.

**Figure 3.**
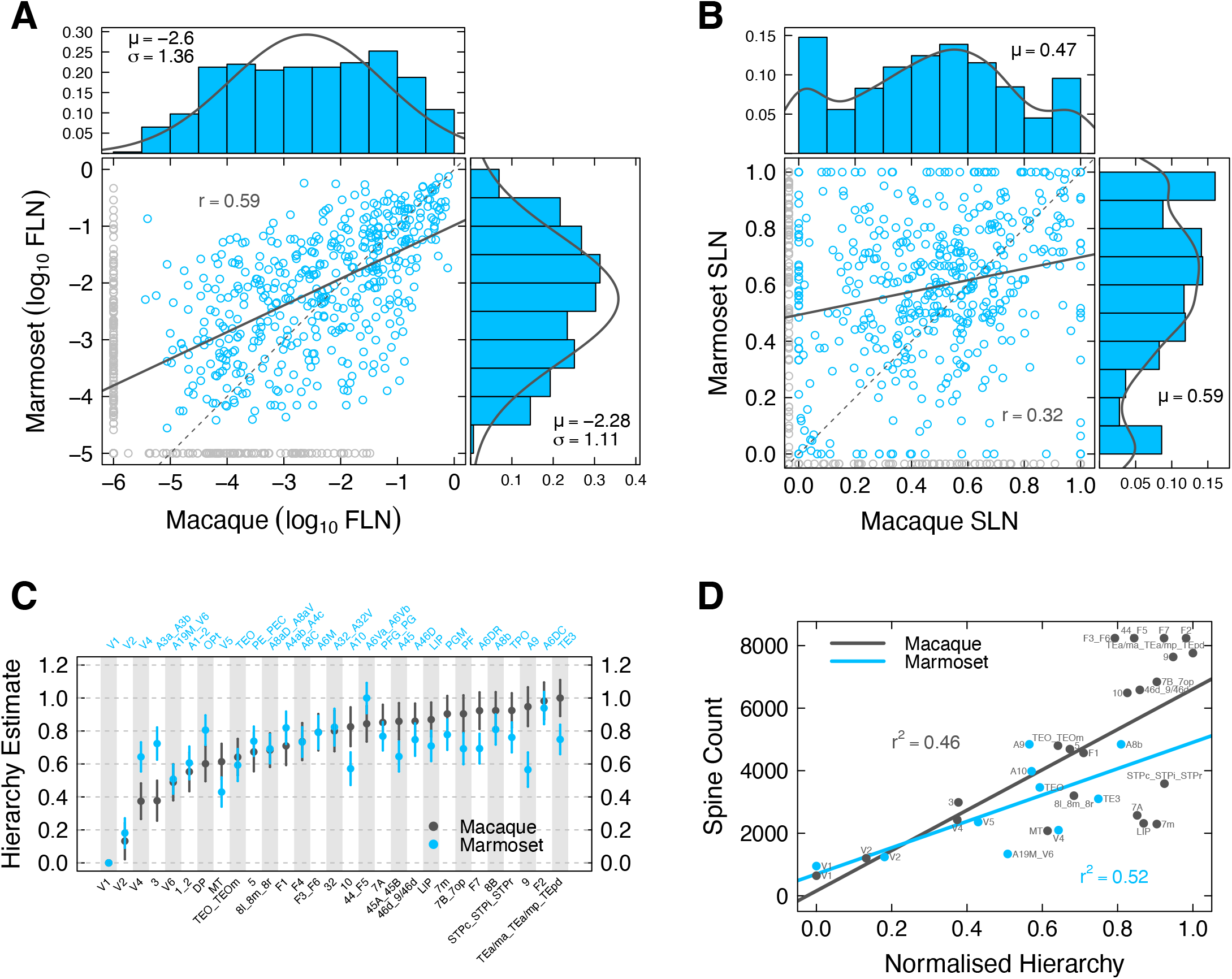
29×29 Connectome statistics, SLN-based hierarchy estimates and spine counts. (A) Macaque (horizontal axis) vs marmoset (vertical axis) consensus log_10_ *FLN* correlation and distribution. In the main plot, each circle represent a connection. Blue, the connection exists in both species; grey, the connection exists in only one species. Solid black line, linear regression line between macaque and marmoset log_10_ *FLN* s, excluding grey datapoints; correlation coefficient of the regression line *r* = 0.59; thin broken line, line of unit slope for visual reference. In the top plot is the log_10_ *FLN* distribution for the macaque, including connections unique to the macaque; mean *µ* = −2.6 and standard deviation *σ* = 1.36, used as parameters for the superim-posed black normal curve. The same statistics are shown in the left plot for the marmoset, the same conventions apply. (B) Macaque (horizontal axis) vs marmoset (vertical axis) consensus SLN correlation and distribution. Same reading conventions as in (A). (C) SLN-based hierarchical estimates for macaque (black dots) and marmoset (blue dots), computed by a GLM of the beta-binomial family, using a logit link function. Values are ordered by macaque increasing estimates. Bottom axis, macaque consensus areas; top axis, marmoset homolog consensus areas. (D) Spine count as a function of normalized SLN-based hierarchy. Black dots and regression line, macaque; blue dots and regression line, marmoset. Proportion of variance explained *r*^2^ is 0.46 for the macaque and 0.52 for the marmoset.

### Macaque connectivity is on average more feedback while marmoset somato motor areas are higher up in the cortical hierarchy

Differences are greater in terms of SLN (Supragranular Labeled Neuron; **Figure 3B**), where correlation between the two species is weak (*r* = 0.32) and where the macaque average (*µ* = 0.47) is lower than that of marmoset (*µ* = 0.59), meaning that macaque connections tend to be, on average, more feedback (FB; originating in infragranular layers). By opposition, the marmoset is tendentially more feedforward (FF; originating in supragranular layers) (**Figure S5**), consistently with the non consensus original data^8,9^. **Figure 3C** shows the difference in SLN-based hierarchical estimates^15,51^ for the two species, with areas ordered on the macaque hierarchy. Marmoset consensus areas A3a A3b (primary somatosensory for touch and proprioception), A4ab A4c (primary motor or M1) and, to a lesser extent marmoset area A1-2 (primary somatosensory for texture, size and shape) and PE PEC (somatosensory dedicated to forelimb digits and joints), which also receives direct somatosensory thalamic inputs^52^, have a higher hierarchical value than their macaque counterparts (macaque areas 3, F1 1 2 and 5 respectively). This, again, is consistent with the pre-consensual data^8^. In other words, somatosensory and primary motor modalities are higher in the global cortical hierarchy in the marmoset than they are in the macaque. At the top of the marmoset hierarchy is consensus area A6Va A6vb (macaque consensus area 44 F5, motor oro-facial homolog to Wernicke in humans).

Conversely, marmoset prefrontal areas A45, A9, A47 (macaque area 12) and A10 have lower hierarchical values than their macaque counterparts. Other interesting differences also occur in the visual system, such as marmoset areas V4, OPt and MIP (V4, DP and MIP in the macaque), which also display higher hierarchical estimates. Thus, a general picture seems to emerge, whereby sensory-motor areas are tendentially higher in the hierarchy for the marmoset, when more associative areas tend to be higher in macaques.

### Marmosets show lower spine count values compared to macaque

Spine counts have been shown elsewhere to correlate positively with SLN-based hierarchy in macaques^31,53^. Here, we show that the marmoset species also displays a correlation of spine count with hierarchy, albeit different in slope (**Figure 3D**). This is yet again consistent with pre-consensus data^8^, thus further validating that the consensus process conserves properties well. In the current consensus data, the maximum spine count density value is just short of 5000, whereas macaque maximal values go beyond 8000. This may be due to lack of data, but avaiable data for the homolog areas in both species suggests a genuine difference in spine counts (see marmoset areas A9, A10, TEO and TE3, and macaque counterparts 9, 10, TEO TEOm and TEa/ma TEa/mp TEpd.

### Macaque and marmoset have different optimal global couplings

Because the 11×11 model does not contain primary sensory areas, both cues (always in excitatory population A) and distractors (always in excitatory population B; **Figure 4D**) are sent to all parietal areas, so as to reflect the distributed input those areas should normally receive from the early visual system. Additionally, sending the cue to a single parietal area fails to create persistent and distributed activity in the system, and only a combination of 4 to 5 parietal areas (depending on elected areas) achieves the goal of distributed sustained activity. Cues are always sent at 1 s into the simulation, and distractors at 4.5 s. Finally, the system is symmetric in the sense that population A and B stable points are identical in terms of firing rate values, as already established in Mejias & Wang^31^.

**Figure 4.**
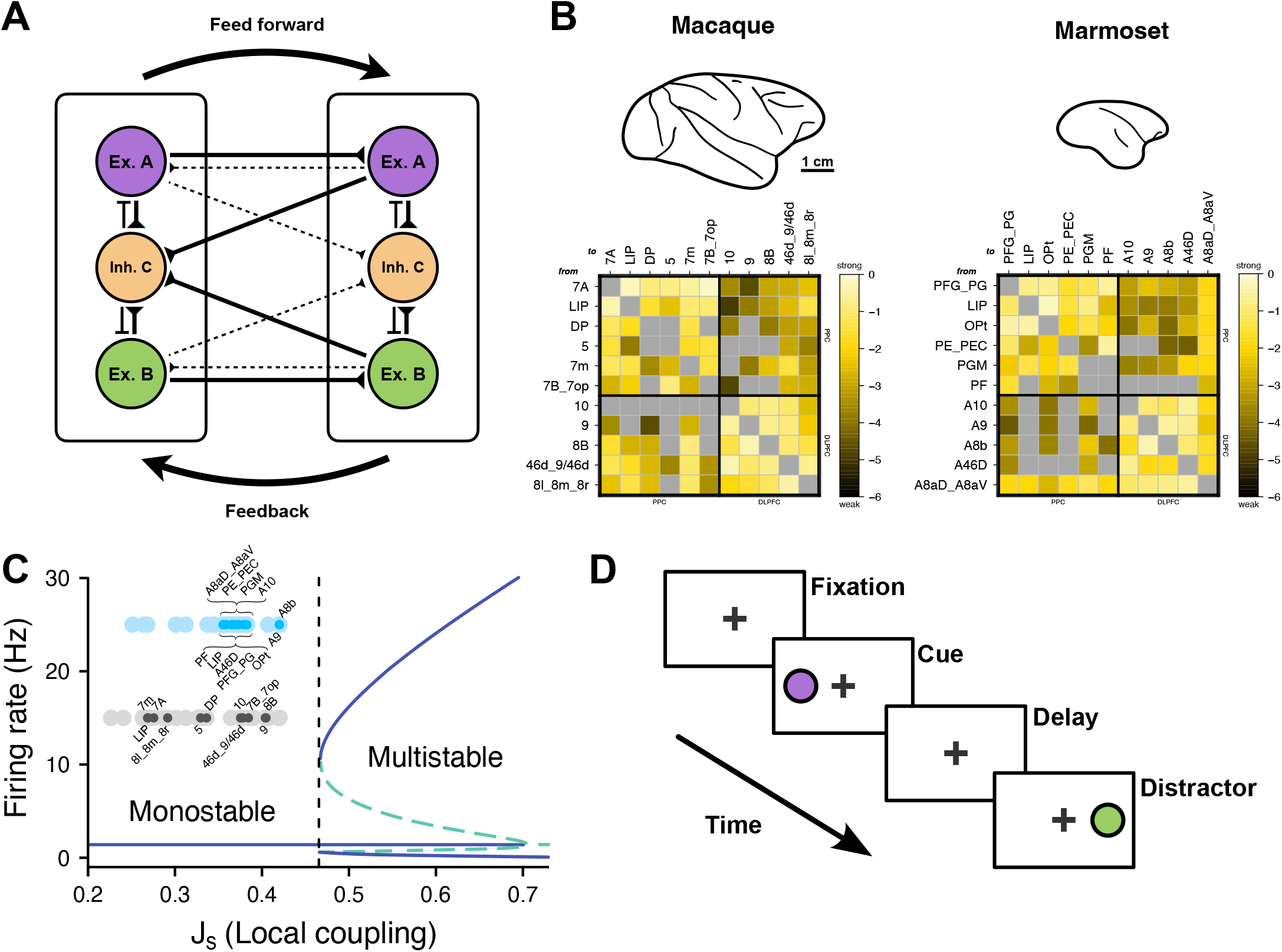
Large Scale model of distributed working memory. (A) schematic of the local circuit and inter-areal interactions. The local circuitry follows a standard 3-population Wong-&-Wang model^68^, with 2 excitatory populations (ex. A and ex. B, purple and green) and 1 inhibitory population (inh. C, orange). Locally, populations A and B excites C which inhibits them in return; globally, inter-areal interactions are based on FLN and SLN anatomical data. (B) Comparative visual showing the macaque and marmoset brains at the same scale; scale bar, 1 cm. Under, the 11×11 submatrices that will be used the WM modeling. PPC, Posterior Parietal Cortex; DLPFC, Dorsal Lateral PreFrontal Cortex. Reading conventions identical to figure 2. (C) bifurcation diagram for an isolated area, with firing rates (Hz) as a function of parameter *J*_*s*_. Below a certain threshold value of *J*_*s*_, a single area cannot sustain long term activity and are monostable. Areas are all parametrized to fall below that threshold. Small dark grey dots corresponds to each of the 11 macaque areas from (B) and indicate their particular *J*_*s*_ values, larger pale grey dots represent the distribution of the rest of the 29 common consensus areas. Regular and pal blue dots assume the same function with respect to the marmoset. (D) Schematic representation of the WM task time course. After a 1 second fixation period, the cue is presented for 0.5 s (purple disk), in the form of a pulse of 0.3 nA sent to population A. After a delay period of 3 s, the distractor is presented (green disk) as a pulse to population B identical in time and intensity to the cue.

All free parameters of this model are fixed at the same values for both species except for global coupling strength parameter *G*, which requires specific parametrization for each species to meet certain basic behavioral criteria. These basic criteria are the following: 1) being able to produce distributed and sustained activity; and 2) not to produce spontaneous sustained activity if not solicited by a cue in either excitatory population.

We determined the valid range of *G* values by doing a systematic parameter search with gradual incrementation of step size 0.01 for both species, using 10 s simulations over 20 different random seeds (**Figure 5A**). Valid range of value go from 0.80 to 1.16 in the macaque, versus 0.57 to 1.13 in the marmoset (averaged over the 20 seeds for both species). Final values of *G* were chosen as the mid point of the resilient range for the macaque and the distractible range for the marmoset, at 0.98 and 0.85 respectively (**Figure 5A**). Specific to the macaque system is the existence of a regime of partially resilient behavior (from 0.45 to 0.80), where prefrontal areas show persistent activity but parietal areas fail to do so (**Figure 5B**, complete set in **Figure S6A**). Additionally, there exists a small, not robust, window of low *G* values (between 0.50 and 0.57) that produces a resilient marmoset (**Figure 5C**, complete set in **Figure S6B**).

**Figure 5.**
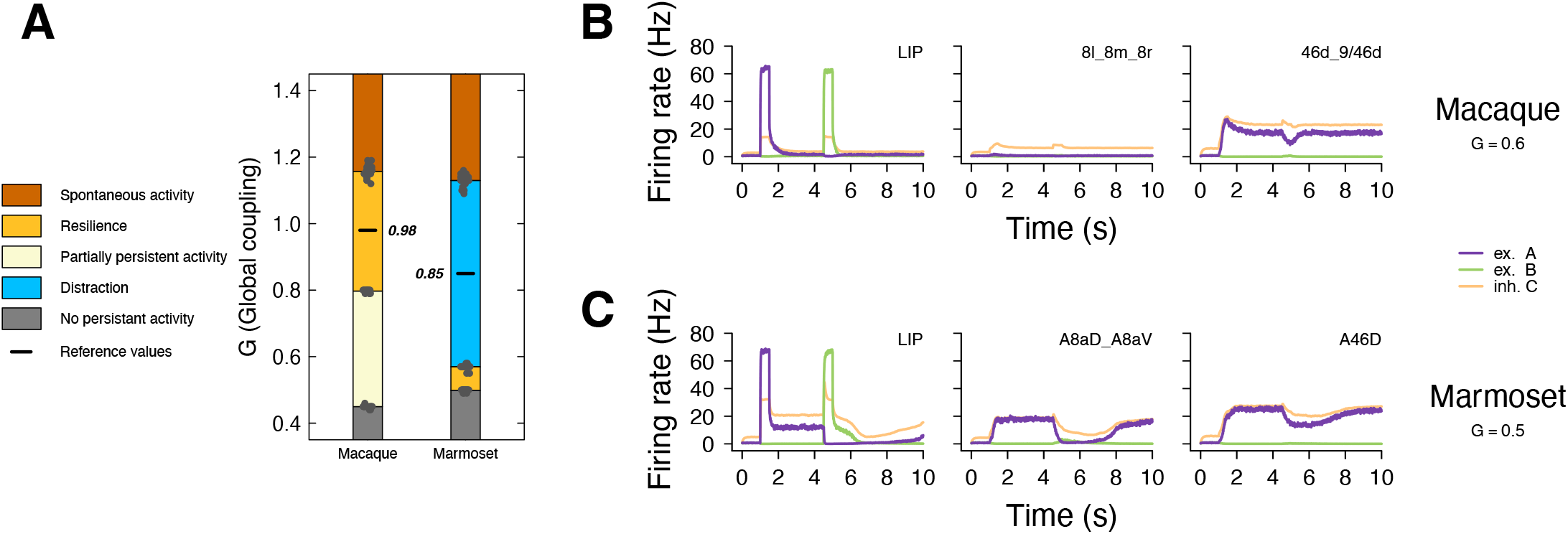
Parametrization of global coupling parameter *G*. (A) Stacked histogram showing different regimes of activity as a function of *G*, for both species. Dark orange, spontaneous activity; yellow, resilience to distraction; pale yellow, partially persistent activity; blue, distraction; dark grey, no persistent activity, black little, reference (chosen) value. Simulations were run with incremental *G* steps 0.01, with 20 different random seeds (black dots). Boundaries between regimes are computed as averages over all seeds at each qualitative change of regime. Reference value for each species is defined as the half point in the resilience regime for the macaque and the distracted regime for the marmoset. (B) Firing rate as a function of time for the macaque’s partially persistent regime at *G* = 0.6. Purple trace, excitatory population A; green trace, excitatory population B; orange trace, inhibitory population C. Areas have been selectively chosen to showcase the partialness of this regime, with parietal areas not sustaining activity (LIP) whilst prefrontal areas do (46d 9/46d). Consensus area 8l 8m 8r (FEF) is shown as the one prefrontal area not following this logic. (C) Example behavior for the marmoset’s small window of resilient regime at *G* = 0.5, where population A re-establishes itself after the distractor pulse. Homolog consensus areas and reading conventions as in (B). Complete set for (B) and (C) available in **Figure S6A and B**.

### The macaque is resilient to distraction, not the marmoset

With *G* now fixed, both species in the 11×11 model display similar behavior in response to the cue, in that both achieve distributed sustained activity throughout the system prior to the onset of the distractor (**Figure 6**, purple trace), as is expected from a number of previous works^27–30,34,45,46^. Note the very low rate of areas DP and 8l 8m 8r in the macaque compared to their marmoset equivalents, OPt and A8aD A8aV. A major difference in behavior arises after sending the distractor in excitatory population B. At that point, the macaque shows a resilience to distraction (**Figure 6A**, purple trace) after a brief transient collapse of cue encoding in PPC (**Figure 6A**, green trace), consistent with single-cell LIP^37,54^ and VIP^55^ recordings. additonally, PFC areas are able to maintain their activity in spite of the distractor affecting PPC areas, encapsulating known results^34,38,39^. In opposition to this, the marmoset’s population B takes over (**Figure 6B**, green trace), thus predicting that the marmoset should be more distractible as distractor encoding activity would propagate to DLPFC. Beyond the susceptibility to distraction shown in **Figure 6**, macaque area DP and consensus area 8l 8m 8r (*i.e*. the FEF), fail to achieve strong activity, contrary to their marmoset counterparts OPt and A8aD A8aV (marmoset FEF). This difference is not dependent on parametrization.

**Figure 6.**
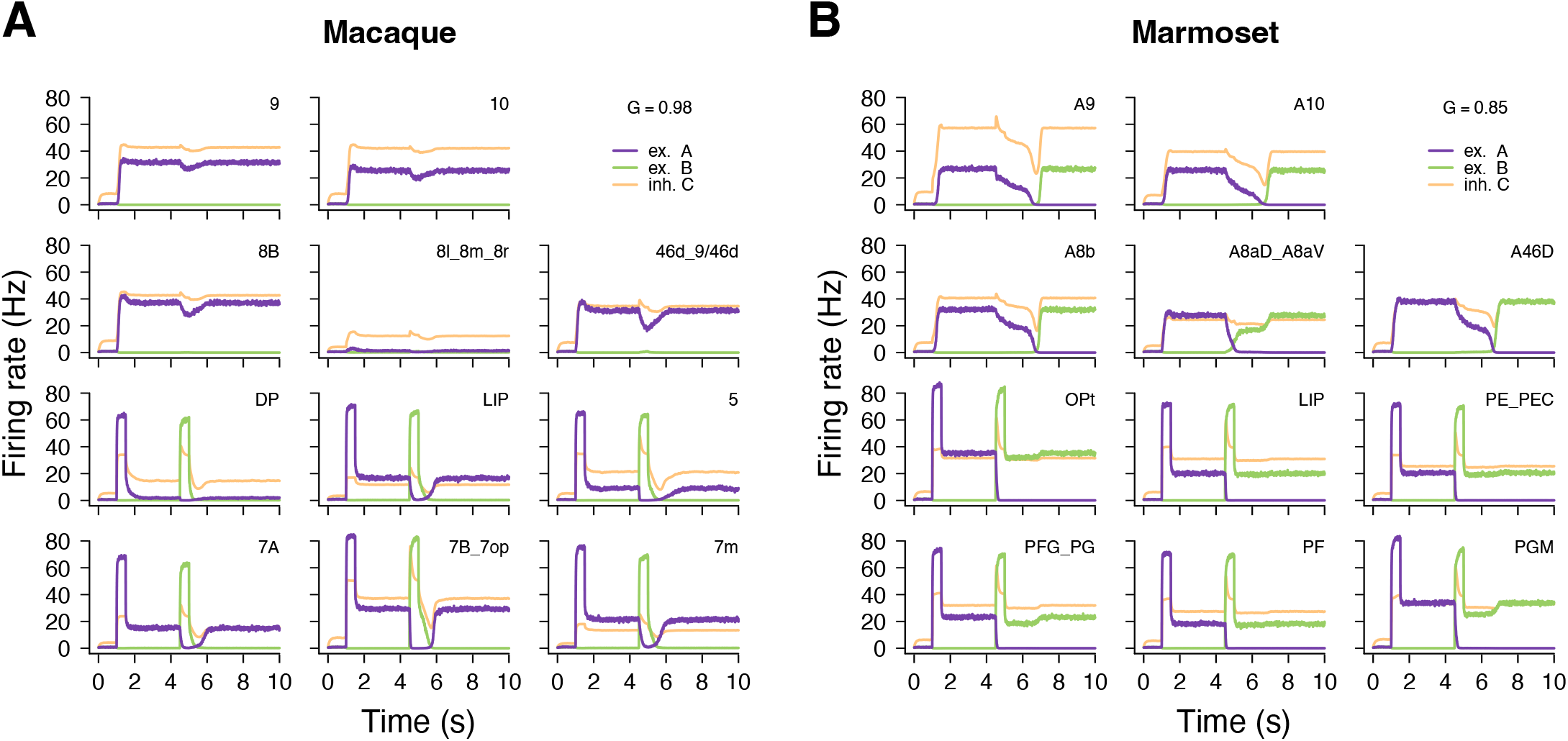
Standard behavior of the distributed frontoparietal working memory model. Each subplot shows the activity traces of the firing rates as a function of time for each of the 11 consensus areas of the selected frontoparietal network. Areas are directly comparable with their homolog and are placed accordingly. Cues (*t* = 1 s to population A, purple trace) and distractors (*t* = 4.5 s to population B, green trace) are sent to all parietal areas. Orange trace is inhibitory population C. (A) Macaque; global coupling parameter *G* = 0.98, resilient regime. (B) Marmoset; *G* = 0.85, distractible regime.

The activity of macaque DP and FEF is not exactly 0, since DP averages around 1.8 Hz and FEF around 1.3. Such a reduced, yet non-zero activity is indication of reduced input from the rest of the network. Looking back at **Figure 3B**, the average of (non logged) FLN input from the parietal is ∼ 0.008 for the macaque and ∼ 0.014 for the marmoset. Cumulatively, the macaque FEF receives from parietal areas less than 0.6 times what the marmoset FEF receives from the same areas (∼ 0.049 vs ∼ 0.082). In comparison, the intrafrontal input to the FEF (*i.e*. from the other prefrontal areas) is greater in the macaque than in marmoset, with non-log averages of ∼ 0.066 and ∼ 0.015 respectively. The cumulative sums of FLNs for each are ∼ 0.263 and ∼ 0.059, that is a 4.4 times larger input for the macaque.

A similar observation can be made for DP/OPt: the connection average FLN from other parietal areas is ∼ 0.061 for the macaque and ∼ 0.153 for the marmoset, for a cumulative input of ∼ 0.183 vs. ∼ 0.765, about 4.2 stronger in the marmoset. comparatively, input from the prefrontal areas to DP/OPt are very weak, averaging ∼ 0.0017 for the macaque and ∼ 0.0006 for the marmoset, with a cumulative sum of ∼ 0.0067 and ∼ 0.0026 (∼ 2.5 higher for the macaque, but still a very weak connection).

Additionally, SLNs are also at play: FEF and DP in the macaque average at ∼ 0.360 and ∼ 0.428 respectively, vs. ∼ 0.698 and ∼ 0.669 for the marmoset. This means that, all things being otherwise equal, FEF and DP are non negligibly more FB in the macaque than they are in the marmoset, which is not the case for the any other area, with the exception of 7A/PFG PG.

In summary, the macaque FEF indeed receives less cumulative parietal inputs and more prefrontal inputs than it does in the marmoset, as is the case for macaque area DP compared to its marmoset homolog OPt. Adding to this the fact that connections to both macaque FEF and DP are more FB than in the marmoset, and therefore more inhibitory *via* the Counter Inhibitory Bias (CIB; Mejias & Wang^31^), we see that the anatomy easily explains the dynamical difference between the two species.

Additional simulations were run with a 2×2 PFC-PPC model on the one hand, and a full 29×29 areas on the other. The 2×2 model shows a similar tendency as the 11×11, with the macaque being resilient and marmoset being partially (but not completely) distractible (**Figure S7A and B**). The 29×29 version of the model fails to yield the same species difference(**Figure S7D and E**).

### Changes in regional coupling can reverse resilience and sensitivity

In order to understand what makes macaque and marmoset different in their distractibility, we investigated how we could make the macaque model susceptible to distraction and, vice-versa, force the marmoset to be resilient to distractors. Swapping around the specifically macaque connections to the marmoset and vice-versa failed to yield the expected result (**Figure S8**). We therefore elected to divide the 11×11 connectivity matrices of both species into 4 quadrants, akin to intra-regional and inter-regional subsets of connections, and attributed to each a specific multiplicative parameter, from *ρ*_1_ to *ρ*_4_ (**Figure 7A**). The parameter space of *ρ*_1_ vs. *ρ*_2_ is showed in **Figure 7B**. Conversion can be achieved in the macaque by reducing *ρ*_1_ to any value below 0.6 (**Figure 7A**; also true for the 2×2 PFC-PPC macaque model, see **Figure S7C**), thus effectively tuning down the intra-frontal connectivity. The consequence of this is to shift the system to a part of the parameter space where distraction can occur (**Figure 7B**, left panel, leftward arrow). Sensitivity to distraction in the macaque can be observed in example areas DP, LIP, 8l 8m 8r (FEF) and 46d 9/46d **Figure 7C**. LIP and 46d 9/46d are shown as good prototypical areas for parietal and prefrontal regions, respectively. DP and the FEF are shown because of their lack of activity in the macaque, as already visible in **Figure 6** (complete set in **Figure S9A**).

**Figure 7.**
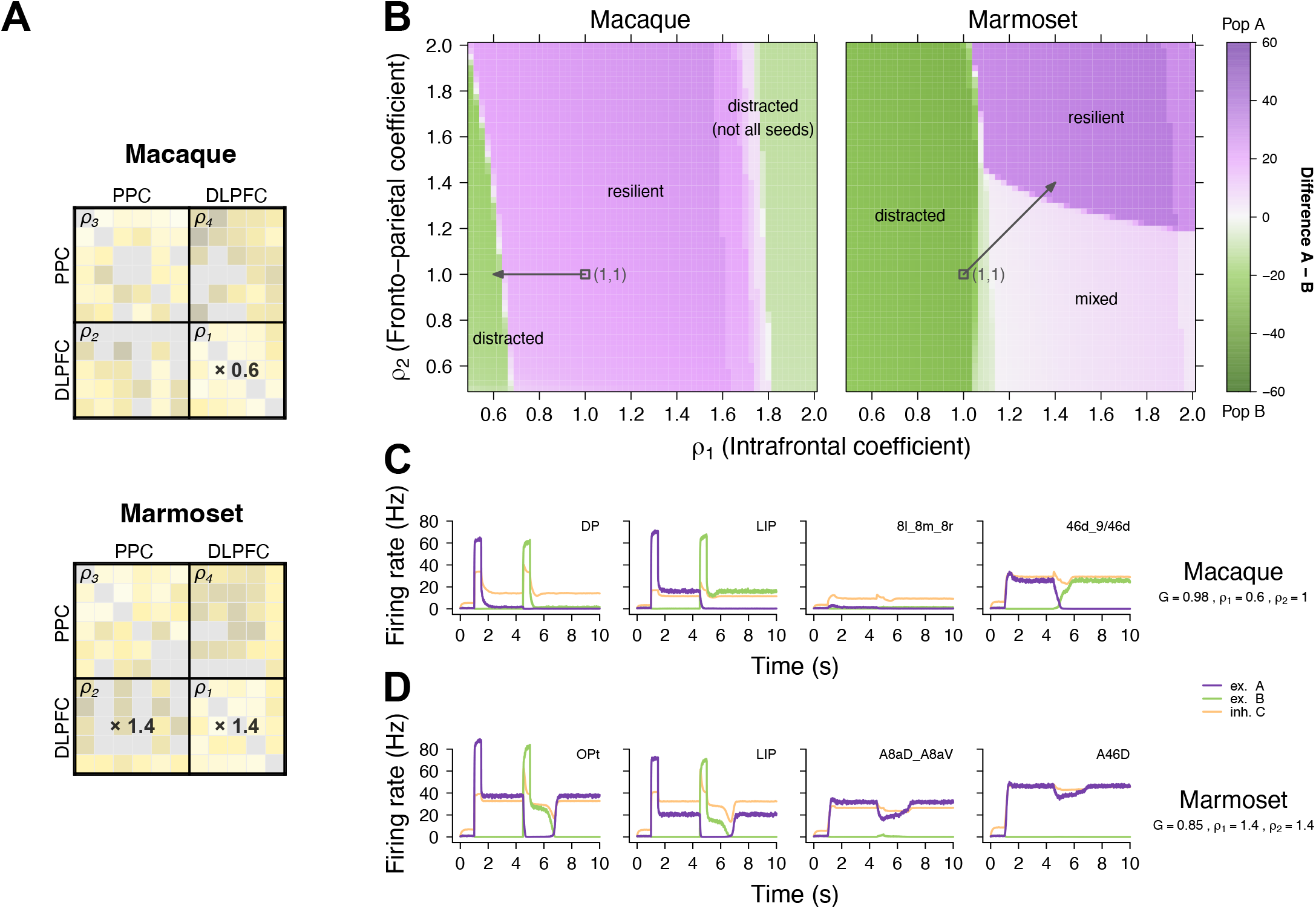
Parameter space analysis of resilience and distractibility. (A) 11×11 matrices from **Figure 3B** divided in four quadrants: *ρ*_1_, DLPFC to DLPFC or intrafrontal; *ρ*_2_, DLPFC to PPC or frontoparietal; *ρ*_3_, PPC to PPC or intraparietal; *ρ*_4_, PPC to DLPFC or parietofrontal. Setting *ρ*_1_ to 0.6 in the macaque makes it sensitive to distraction. Setting *ρ*_1_ and *ρ*_2_ to 1.4 in the marmoset makes the system resilient. (B) *ρ*_1_-*ρ*_2_ parameter plane for both macaque (left panel) and marmoset (right panel). Green hues indicate resilient territories in the plane, purple hues indicate distracted, and white indicates mixed behavior (see **Figure S6C**). Each square is the average of simulations performed over 10 different random seeds, by incremental steps of 0.025 in both directions. Black open squares indicate (1,1) coordinates in the plane, black arrows indicate the displacements generated by values in (A). (C) Example behavior for the macaque’s distractible regime at *G* = 0.98, *ρ*_1_ = 0.6, *ρ*_2_ = 1. (D) Example behavior for the marmoset’s resilient regime at *G* = 0.85, *ρ*_1_ = 1.4, *ρ*_2_ = 1.4. Reading conventions as in **Figure 5B and C**. Complete set for (C) and (D) available in **Figure S9A and B**.

As can be seen in **Figure 7B**, the macaque possesses what appears to be another distracted territory for very high values of *ρ*_1_, with gradual transition roughly centered around 1.8. However, this part of the parameter space is in fact unstable as the outcome behavior depends on the random seed used in any particular simulation. This is indicative of a parameter going beyond its “physiological”, or at least realistic, limits. At these levels, any random fluctuation is amplified by a factor of close to 2 in the prefrontal areas. Helped by a higher firing rate baseline also due to the higher *ρ*_1_ value, this leads to an easy propagation to rest of the network. Due to their unrealistic nature, these will not be discussed further in what follows.

A marmoset resilient to distraction can similarly also be achieved by giving *ρ*_1_ and *ρ*_2_ the value of at least 1.4 each (**Figure 7C**), which effectively displaces the system to a part of the parameter space that grants it resilience (**Figure 7B**, right panel, right-upward arrow), with example areas showcasing the behavior in **Figure 7C** (Complete set in **Figure S9B**). A mixed behavior regime exists in the marmoset’s *ρ*_1_-*ρ*_2_ plane, which is characterized by partial activations in the system (**Figure S9C**). The 2×2 PFC-PPC marmoset model effectively sits in this regime. Consistently, a single change of *ρ*_2_ to 1.7 pushes the 2×2 system upward into resilient territory (**Figure S7C**).

As a side note, similar parameter spaces can be achieved with *ρ*_3_ and *ρ*_4_ that also show different domains of resilience and susceptibility, which are topologically similar to the ones of *ρ*_1_ and *ρ*_2_ (**Figure S10C**). There, the macaque becomes sensitive to distraction as one increases *ρ*_4_, and the marmoset becomes resilient by reducing both *ρ*_3_ and *ρ*_4_. For practical purposes, the focus will bear on *ρ*_1_ and *ρ*_2_ in what follows.

### Gradual changes in regional coupling indicate sharp transition between resilient and distracted regimes

The next logical step was then to try and understand the transition from one regime to the other. To do so, we performed simulations at incremental steps in the *ρ*_1_ parameter space dimension, across the border that separates the two regime, and looked at the conductance *S*_*A*_-*S*_*B*_ space (*i.e*. Excitatory populations A and B’s *S* variable) for the system’s trajectories, using each time 10 different random seeds. For the macaque, *ρ*_2_ was fixed at a value of 1, and *ρ*_1_varying from 0.75 to 0.55 (*i.e*. from resilient to distractible) with steps of 0.01 (**Figure 8A**, *ρ*_1_-*ρ*_2_ panel). **Figure 8A** displays the state of the macaque *S*_*A*_-*S*_*B*_ phase portrait (rotated by *π/*4) for increasing values of *ρ*_1_. At *ρ*_1_ = 0.72 (first panel on the left, full progression visible in **Video S1**), the macaque displays its standard behavior. The at-rest stable fixed point (black dot) is reached within a small number of time steps (black trajectory originating from coordinate (0,0)) and the system will stay there until the cue is delivered. Said cue can be sent to either population A (purple trajectory) or population B (green trajectory). In either case, the system will reach the stable fixed point specific to the population that was cued (purple point for population A, green point for B), after a quick overshoot. At that point, a distractor can be injected in the opposite population. When doing so, the system’s trajectory shoots inward for the duration of the pulse (trajectories yellow and red), only to come back the same stable fixed point they were at before distraction.

**Figure 8.**
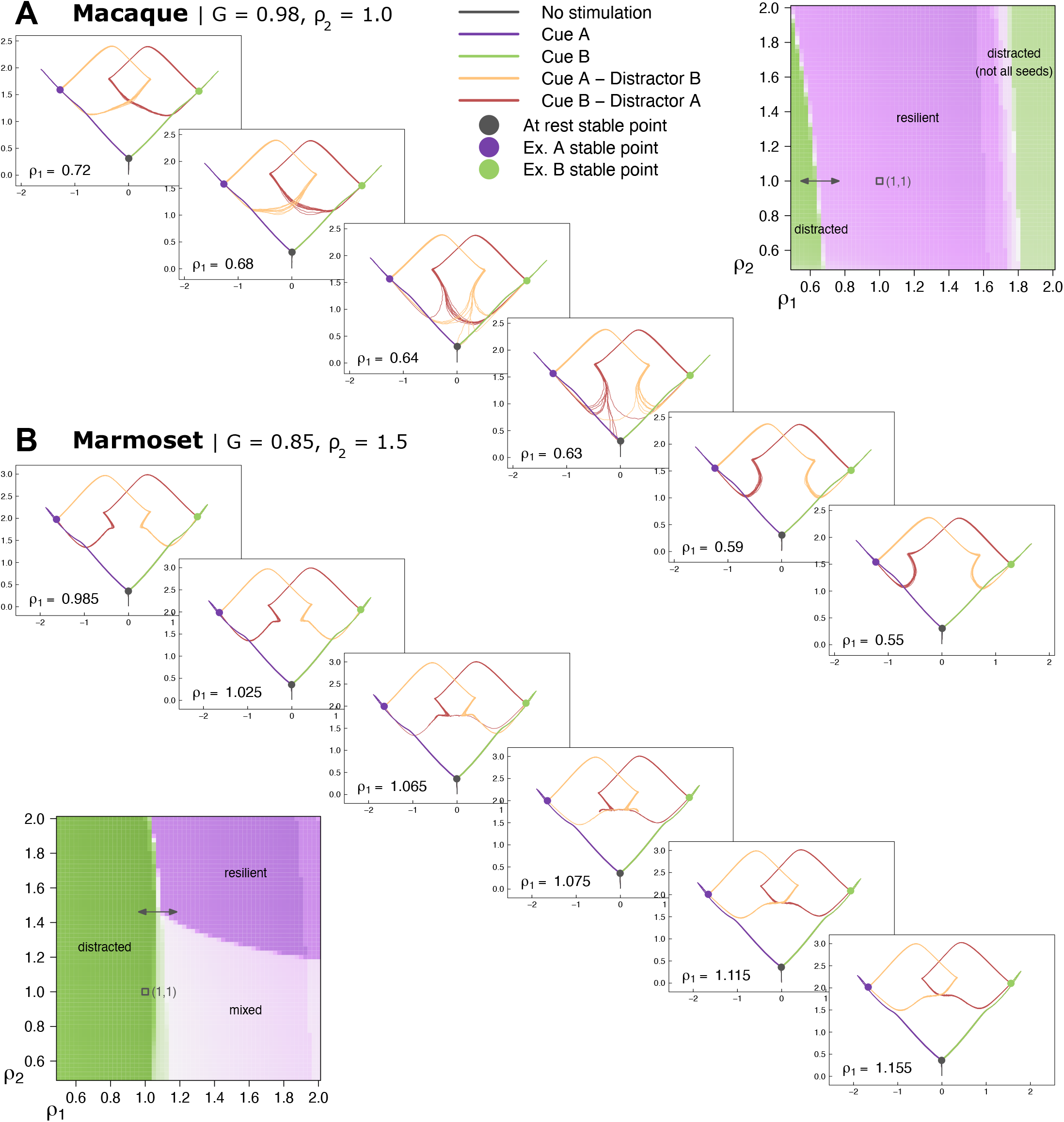
*S*_*A*_-*S*_*B*_ conductance phase space gradual change across regime boundaries. The plane is chosen as the unique plane defined by the 3 stable fixed points of the system, then rotated by *π/*4 for visual clarity. Colored dots indicate said fixed points: black, at rest (no input current); purple, population A; green, population B. Colored lines follow the same convention for trajectories: black, at rest (no input current); purple, cue input current in population A; green, cue input current in population B; yellow, distractor input current in population B when at stable fixed point A; dark red, distractor input current in population A when at stable fixed point B. Trajectories computed for 10 different random seeds for all conditions. (A) Macaque. Each plot corresponds to an incremental step in *ρ*_1_, starting from a resilient macaque. Parameters *G* and *ρ*_2_ are fixed at 0.98 and 1 respectively. Full investigated range from *ρ*_1_ = 0.75 to *ρ*_1_ = 0.55 with 0.01 step size, available in **Video S1A**. (B) Marmoset. Each plot corresponds to an incremental step in *ρ*_1_, starting from a distractible marmoset. Parameters *G* and *ρ*_2_ are fixed at 0.85 and 1.5 respectively. Full investigated range from *ρ*_1_ = 0.975 to *ρ*_1_ = 1.175 with 0.01 step size, available in **Video S1B**. For each species, inset plots help visualize the covered range of values in *ρ*_1_-*ρ*_2_ parameter space.

Trajectories will become increasingly tempted by the other side as *ρ*_1_ decreases. Transition into to the distractible regime of behavior for the macaque occurs over a very narrow band of *ρ*_1_ values, contained between 0.64 and 0.63, where independent random seeds show radically different trajectories, some showing resilience and returning to their fixed point of origin, and others showing susceptibility and ending their course on the opposite stable fixed point. From *ρ*_1_ = 0.62 downward, the system has fully transitioned to being distractible and all seeds display similar behavior once again, this time to be diverted to the opposite fixed point.

Looking now at the marmoset, we followed an identical procedure, only fixating *ρ*_2_ at a value of 1.5 so as to avoid going through the mixed regime of behavior (**Figure 8B**, *ρ*_1_-*ρ*_2_ panel). *ρ*_1_ was there set to vary this time from 0.975 to 1.175, again with steps size of 0.01. Qualitatively, the behavior is reversed from the macaque, as the marmoset is by default sensitive to distraction. At *ρ*_1_ = 0.985, well before the transition happens, trajectories are quite analog with those of the distractible macaque. As *ρ*_1_ progresses toward the threshold of behavioral shift, random seed trajectories remain well grouped together, and will remain so during and after the transition, which is an interesting qualitative difference with the macaque, arguing for a qualitatively different high dimensional landscape, as they might in fact be fanning out in a different plane. The transition itself even shorter than in the macaque, as all seeds have transitioned at *ρ*_1_ = 1.075 to a fully resilient marmoset. Interestingly, these distractor-based trajectories still show a small detour along their former distractor sensitive path before turning around toward their newly found target. The last of this can be seen at *ρ*_1_ = 1.105 (**Video S2**).

## DISCUSSION

### Consensus mapping conserves statistics and is a great tool for future comparative work

The consensus mapping that we have exposed and successfully used here offers the unique opportunity to directly compare the mesoscale connectomes of two species of primates. Allowed by the extensive anatomical analogies that exist between primates, the consensus mapping is largely based on the merging of subareas together to recover coarser-grained areas. This comes at the price of loosing some of the spatial granularity for either species. Nonetheless, as we have seen with **Figure 3 and S5**, the consensus process conserves the basic statistics, FB and FF distributions, graph densities and spine count to hierarchy relationship^8,18,21^ of each species pre-consensus. Therefore, the mapping is stable enough to be meaningful for the modeling depicted in this study. Future anatomical investigations of these two species will continue to improve the overlap of injected areas, and should prove important for future comparative work in primate anatomy.

### Distractibility is intrinsic to the marmoset model

The most crucial finding of this study is the fact that the macaque and marmoset models, which are identical except for their anatomical data, capture an important and real cognitive difference between the actual species in their resilience to distraction. If this result is consistent with known macaque capabilities and electrophysiology^27–30,34^, marmoset research is still at too early a stage to confirm our prediction, although available results certainly point in the same direction^45,46,49^.

For the readers to assess carefully the validity if this result, some aspects of the modeling need to be further discussed. The first of them is how the global coupling parameter *G* (**Figure 5A**) value was chosen. In the macaque, the parametrization is fairly straightforward, given that there is only one bracket of values that produces a fully distributed activity without running the risk of spontaneous activity, with no possible values offering the possibility of a distractible system. The marmoset on the other hand, although its largest window of viable activity is indeed the distractible one, does have a valid, if narrow and less robust, window of resilient behavior. Therefore, purposefully choosing a distractible *G* value to then claim that our model captures differential distractibility may at first glance look like a self-realizing prophecy. However, that would be missing the point, which is that the macaque possesses no distractible *G* values when the marmoset is mostly distractible for the same range of values, thus giving vastly different resilience/distractibility profiles as a function of *G*. Distractibility in the marmoset is therefore a strong prediction of our modeling, softly confirmed by the 2×2 modeling. The only minor difference is that the marmoset defaults there to the mixed behavior regime, understood here as an incomplete distractibility, but still very much closer to it than the macaque. Given that the models are differing by anatomy only, this *G* profile difference can only be due to the variations in anatomy across the species. This means that a network-distributed, complex-system effect from structure to function is here at work, as no single change in the connectivity matrix can produce this behavioral difference on its own. A second aspect of parametrization concerns the parameters *J*_*s*_ and *J*_*IE*_ which, albeit fixed for each area, will vary as a function of the available spine count gradient data.

As shown earlier, the 29×29 version of the marmoset model fails to achieve distractibility (**Figure S7D and E**), and it does beg the question as to why the 11×11 sub network that we use effective captures the dominant mechanism underlying distraction resistance. First, this 11×11 stems from an expansion from Murray et al., 2018^40^ (see Method section: Consensus mapping: 11×11 matrices). Second, there is a growing understanding that different brain states, such as sleep, wakefulness, attention, vision, relaxed mind-wandering, and indeed working memory, have network counterparts of specifically active and synchronized cortical areas^56–58^, whilst others tend to desynchronize for the duration of that state, as seen in state transitions^59^. Selecting for WM network^27,33^ in this study is meant to be an analog of the brain being in that specific state. To recover the result while keeping the full 29×29 network would require modeling state transitions, by implementing a modulatory mechanism that gates out the other areas, potentially based on neurotransmitters such as dopamine^49,60^ or other modulatory systems^32^.

### A larger brain increases feedbackness and therefore stability and resilience to distraction

The very low level of activity in macaque FEF and DP, compared to their marmoset counterparts, can be explained by the difference in connectivity for these areas across the two species. As we have argued elsewhere^19^, these anatomical differences can be thought in the framework of the scaling properties of the brain. Indeed, due to physical constraints exerted onto the brain^61^, a bigger brain will get bigger only the cost of reducing its overall connectivity, which in turns leads to an increasing modularity of its network^19,62^. This effect of scaling the brain up may well be at work here: the macaque and the marmoset 29×29 consensus connectivity matrices in **Figure 2** have a similar graph densities, although a 4% drop is already detectable from marmoset (0.72) to macaque (0.68). Additionally, **Figure 3A** clearly shows that the macaque’s FLN range spans 6 orders of magnitude when the marmoset does only 5^8,14^, meaning that the former possesses extra-weak connections compared to the latter. In short, the macaque brain is already about twice as big as the marmoset brain and, although it is still small enough to harbor a high graph density, it is already on its way to partial disconnection compared to its smaller cousin, as already shown elsewhere^8^.

From a different perspective, Markov et al. ^15^ showed that FB connections decay with white matter distance more slowly than FF ones. This means that, on average, long-range, weak connections will tend to be FB. Extrapolating across species, we should expect the brain to become increasingly FB-dominated as it grows larger, at least in primates. This aligns well with the data shown in **Figure 3B**, where the mean SLN value for the macaque is 0.47, which is non negligibly lower than the 0.59 of the marmoset, meaning that the macaque is on average more FB than the marmoset. Given the implementation of the Counter Inhibitory Bias (CIB; Mejias & Wang^31^) in the system, FB connections target the inhibitory population C, meaning that an on average more FB connectivity will lead to greater global inhibition. This is consistent with the macaque being resilient, as the random distractor will enter a system that is in effect less suited to its propagation. Conversely, the marmoset system is well suited to welcome the next distractor, its global feedforwardness effortlessly carrying its excitatory influence to all areas.

Circling back to macaque FEF and DP, these are arguably the deeper reasons why the macaque FEF and DP fail to display high levels of activity in our simulations. A bigger brain leads to some amount of disconnections — weaker connections or even missing ones — and greater feedbackness, which, in turn, leads to less input — although more selective — for these areas. From this, a conjecture could be that the same model grounded on a bigger brain’s connectivity, such as that of a great ape or even a human primate, would see further disconnections and even greater feedbackness and comparatively more areas behaving in that particular fashion. This, in turn, would be consistent with the increasing well segregated sub-network and functions that we see appear in bigger brains^19,63^.

### Reversing distraction and resilience is consistent with a bifurcation in highl non-linear dynamical systems

As we saw with **Figure 5 and 6**, this system produces by default a macaque resilient to distraction and a marmoset sensitive to it, on par with the behavioral data available at this time^47,49^. The anatomical frontoparietal differences between macaque and marmoset, such as the missing connection between consensus marmoset areas A46D and LIP, known to be highly relevant for WM in the macaque^27^, fail to explain the difference in behavior within the constraints of our model. Even the complete swapping of species specific connections from one to the other cannot achieve the intended behavioral reversal (**Figure S8**). This argues for the existence of an emergent stability produced by network-based complex system of non-linear interactions.

This stable default behavior can nonetheless be changed by regionally (*i.e*. DLPFC vs. PPC) altering the connectivity. The macaque becomes distractible by artificially scaling down the prefrontal self connectivity, coherently with previous work^40^ and effectively making it less powerful at talking to itself and less prone to down-regulating parietal areas. This is consonant with experimental data, where PFC lesions increase distractibility in macaques^34^. On a more graph theoretical side, it can be thought of a way to reduce the prefrontal’s modularity. Because this alone is enough to allow a distractor to win over the system and because the distractor signal comes from the parietal areas, this result can be interpreted as making the prefrontal more permeable to upward signal from the parietal.

In the marmoset’s case, both intrafrontal and frontoparietal connections need to be tuned up to achieve resilience which, at a conceptual level, creates a nice and logical continuum between the two species. However, the opposition is imperfect. Were it to be perfect, an intrafrontal increase would have been enough to render the marmoset resilient. In practice, one needs to increase the frontoparietal connections (*ρ*_2_) by a similar amount to complete the transition, as is shown in **Figure 7B**. Increasing the *ρ*_1_ to 1.4 but keeping *ρ*_2_ at 1 lead to a mixed regime where prefrontal areas are resilient but parietal areas remain sensitive to distractors**Figure S9C**. The likely interpretation is that, by increasing only intrafrontal connections, each prefrontal area now receives enough input from the other prefrontal areas that their collective activity is self-sustained and hard to break. We know from Murray et al. ^40^ that resilience to distraction is given by the prefrontal module. For this new found capability to propagate to the rest of the system, one needs to increase the influence of prefrontal areas on the parietal, which is here modeled by increasing *ρ*_2_.

This need of raising *ρ*_2_ to complete the transition may come from the missing connections between marmoset area 46 (here A46D) and parietal areas LIP, OPt and PE PEC (**Figure 4B**), given the importance of area 46 to WM^27^. Although the marmoset possesses weak frontoparietal connections from A10 to the medial parietal cortex that the macaque does not (**Figure 4B**), these are unlikely to provide robust top-down control. Many substitutions of connections between the two species, analogous to grafting experiments, were tried in the course of this study, but no particular change in the matrix gave a clear cut answer. Incidentally, this is also true with respect to both SLNs and spine count interchanges. all theses are signatures of a complex system.

Now imagining the phase diagram shown in **Figure 8A** as a 3D energy landscape seen from above, becoming distractible here means that the wells that constitute population A and B stable points are getting shallower, with an increasingly reduced basin of attraction that can keep the system captive to that fixed point (*i.e*. the separatrix). As *ρ*_1_ gradually decreases, so does the depth of the wells and the size of the separatrix. Thus, there is a point at which the distractor’s energy is enough to escape (*ρ*_1_ ≈ 0.64), only to fall into the other fixed point’s energy well.

The same is true for the marmoset, only reversed with respect to *ρ*_1_. As the parameter increases (**Figure 8B**), the stable fixed points are gradually pulled deeper, increasing the basin of attraction in both size and gradient, and making its separatrix gradually less escapable. starting at *ρ*_1_ ≈ 1.065, the energy wells are beginning to be too deep and/or too steep for any distractor to be able to push the system out of the well, and the system become resilient to distraction.

As can be seen in **Figure 7B** and further in **Figure 8**, the transition between distracted and resilient regimes is fairly sharp in both species, and leads to radically different qualitative results. In dynamical systems, this corresponds to the general definition of a bifurcation^64^. As it does not appear to depend on local changes in the quality or quantity of stable or unstable fixed points, it should be better seen as a global bifurcation, although there are no limit cycle seemingly involved here. Nonetheless, the conceptual definition does seem to apply, as it appears obvious that some aspects of the high dimensional topology of the phase spaces in **Figure 8** are drastically changing over a very short span of *ρ*_1_ values.

### Distraction may be adaptative given the marmoset’s ecological and ethological niche

From the bifurcation perspective hereabove, it appears that macaques and marmosets effectively sit on opposite sides of a common bifurcation. The particular side on which each species finds itself has been shaped by evolution and should relate to pertinent ecological and ethological aspects of their natural life. As further proof of this, the recent work of Joyce et al. ^49^ have shown that marmoset DLPFC parvalbumin neurons have higher density of D1 (dopamine) receptors, which leads in their modeling to heightened susceptibility to distraction. Further, those levels are increasing from humans to marmosets, which in turn have similar levels to mice.

The first explanation that comes to mind is brain size, as we have made the case elsewhere that it is the prime factor that will allow for the sufficient segregation of functions, itself necessary for higher cognitive capabilities^19^. Further, and not unrelated, we explained earlier how weak, long range connections are tendentially more FB, leading to the expectation that larger brain should also be tendentially more FB, all things being otherwise equal. Another important ecological aspect to consider is their position in the trophic chain of their respective ecosystems.

Indeed, by sheer size effect, the predatory difference may well select for heightened alertness in marmosets, which should in part rely on greater distractibility for either alarm calls from the group or direct sensory contact with a predator.

Finally, this distractibility difference aligns well with the differences at work in their social ethology. Where macaques heavily rely on aggression and distrust^65^, marmosets are a highly collaborative species with respect to problem solving, a trait otherwise fairly restricted to apes^66,67^. In that particular context, distraction may be better seen as the local aspect of a larger, social information flow. Conversely, in a strict hierarchy of aggression, the necessity to efficiently navigate complex power dynamics, an individual macaque able to retain and manipulate information on their own, despite distractors coming from the group, would likely fare better than average.

An in depth research in social cognition, ethology and ecology would be necessary to untangle the complex web of selection pressures that may have shaped differentially our two species, and are far beyond the scope of this particular study. Additionally, Future modeling of the highly non-linear evolutionary dynamics at play here is necessary to deepen or knowledge of these interactions.

## Conclusion

After producing a consensus mapping between macaque and marmoset’s atlases, we reduced each to an homolog parcelation scheme, thus allowing direct, area-to-area comparison of their respective connectomes. We then used these consensus data to differentially constrain an otherwise identical large-scale model of WM. Selecting for the frontoparietal WM network of the same 11 common consensus areas in the two species, we first showed that these two systems produce sustainable activity throughout the network after a cue is sent in. We went on to test their resilience to distraction and discovered that, although the macaque-constrained model resists easily to distraction, the marmoset model readily switches and commits the newest cue to memory by propagating it to all areas, taking over as the new sustained activity. Moreover, the two species can be bridged by scaling up or down specific quadrants of their connectivity matrix, which effectively alters intrafrontal and frontoparietal influence over the system, allowing us to better understand the differences between those two primates. We argue that these behavioral differences between models align intriguingly well with real life behavioral difference, and offer a credible prediction to the marmoset’s behavior in this specific task.

## Limitations of the study

With respect the scaling up of the model to 29 areas (**Figure S7D and E**), the enhanced stability given by the sheer number of areas forbids even the marmoset model to be sensitive to distraction. This points to a limitation of this multiregional model which, due to its architecture, tends to be resilient to distraction already with two areas^40^. The consequence of that is that the marmoset’s 11 area model distractibility is very much non trivial. As scaling up the number of areas tends to lead to resilience from distraction, it means that there is something very specific about the marmoset WM subnetwork’s anatomy that manages to prevent said resilience.

It is only fair to recognize that we have, at this date, less available data for the marmoset (**Table S2**. For non consensus data, see Theodoni et al. ^8^, Supp. Table 4) regarding spine counts than we currently do for the macaque (**Table S1**). Therefore, if the models are indeed identical across our two species, the marmoset may be considered at this point less constrained than the macaque. New data, or data updates that would come in the future may very well change the present results, by making them more alike or more different, by reducing the window of distractibility of *G* or by expanding it. New macaque data could possibly create a band of distractibility on the *G* parameter line. This point, incidentally, is just as true for any change in the connectivity.

Furthermore, as compelling as the proposed hypothetical link between anatomy and behavior may be, it cannot possibly be the entire story. Indeed, many other explanatory factors are by necessity at work in the production of this behavioral difference, from microcircuits to other anatomical gradients, neuronal subtypes, eletrophysiology, etc. What this study shows is that the differential anatomy between those two species, at least in the subnetwork of WM, does predict some behavioral difference with respect to the presence of distractors, which appears to be coherent with the behavioral differences so far known to experimentalists working on these species. Future work in large-scale modeling, as well as modeling of evolutionary dynamics, will need to incorporate increasingly finer-grained details of neurophysiology in order to fully disentangle the links between structure and function.

## RESOURCE AVAILABILITY

### Lead contact

Requests for further information and resources should be directed to and will be fulfilled by the lead contact, Xiao-Jing Wang.

### Materials availability

This study did not generate new materials.

### Data and code availability

1. Complete consensus meso-connectome data, original code and supplementary videos S1 and S2 has been deposited at Zenodo under the DOI 10.5281/zenodo.15612783 and is publicly available as of the date of publication.
2. Any additional information required to reanalyze the data reported in this paper is available from the lead contact upon request.

## ACKNOWLEDGMENTS

The Authors would like to thank Jorge Mejias for providing the base code and helpful advices along the way, the Wang lab for the constant challenges and discussions that helped build this paper, in particular Aldo Battista and Wayne Soo for so gracefully teaching their impressive mathematical knowledge. We would also like to thank Seán Froudist-Walsh and Mary Kate P. Joyce for the helpful discussions on the marmoset neurophysiology and behavior. This work was made possible thanks to the NSF NeuroNex grant 2015276 to A.F.T.A. and X-J.W.; the NIH grant R01MH062349 to X-J.W.; the Simons Foundation grant 543057SPI to X-J.W.; the National Health and Medical Research Council grant APP1194206 to M.G.P.R; the Australian Research Council grant DP210101042 to M.G.P.R; and the Australian Research Council Grant DP210103865 to M.G.P.R.

## AUTHOR CONTRIBUTIONS

Conceptualization, L.M., P.T., M.G.P.R, X-J.W. ; methodology, L.M., P.T., M.G.P.R, X-J.W.; software, L.M.; validation, L.M., P.T., M.G.P.R, X-J.W.; formal analysis, L.M.; investigation, L.M., P.T., A.F.T.A., X-J.W.; resources, L.M., M.G.P.R., X-J.W.; writing – original draft, L.M.; writing – review & editing, all authors; visualization, L.M., P.T.; supervision, X-J.W; funding acquisition, A.F.T.A., M.G.P.R., X-J.W.

## SUPPLEMENTAL INFORMATION INDEX

Figures S1-S10 and their legends in a PDF

Table S5. Complete consensus meso-connectome data organized as a table in an Excel file, too large to appear in the PDF

Video S1 and S2. bifurcation videos related to Figure 8

## STAR METHODS

### Key resources table

### Method details

#### Anatomical connectivity data

Despite the continuous progress occurring in the field of diffusion MRI and tractography based connectomics^69,70^, it nonetheless remains true that, where feasible, retrograde tract-tracing experiments still constitute the gold standard in cortical connectivity/mesoscale connectomics^71^. In the realm of primates, we are now uniquely suited to tackle cross-species comparison as extensive databases have been produced in both the macaque^14,60^ downloadable from core-nets.org, the historical model in primate neuroscience, and now the marmoset^8,9^ downloadable from analytics.marmosetbrain.org, a smaller primate model that is easier to handle whilst retaining many primate features important to today’s neuroscience.

Injections of retrograde tracers are carried out in multiple areas, designated as “target” areas, over multiple experiments. Each injection site is controlled so that the injection covers all 6 layers of the cortex, whilst not infringing on the white matter, so that the dye is not captured by fibers of passage. At the injection site, the dye is captured by axon terminals and is actively transported backward toward the soma of the cell, thus only labeling neurons that directly project onto the injection site (**Figure S1A**). These neurons are subsequently detectable through fluorescent microscopy. Tracers here employed are monosynaptic, meaning that they cannot continue their course beyond the one neuron that captured them into another neuron that shares a synapse with the back-labeled cell.

Many of the injected areas in both species are represented only by a single injection. It has been shown that the connectivity patterns of repeated injections are remarkably stable across individuals for injections in V1, V2 and V4 for retinotopically equivalent points in each of those areas^50^ (in this case foveal). The same observation had already been made for marmoset visual area MT^72^. Importantly, in both species this repeatability includes weaker projections previously unknown to the field. The same analysis was repeated in frontopolar area 10^14^, on the opposite side of the brain, and in the motor cortex^73,74^, thereby proving that this repeatability is not a feature restricted to the visual system. This statistical stability erodes as the parcellation scheme becomes finer and finer grained, and becomes stronger with a coarser parcellation^14^, such as our consensus mapping. This means that a single properly done injection is statistically sufficient to get an accurate and stable connectivity pattern, with repetitions providing only limited additional information. This is also reported to be true for mouse^13^ and the marmoset^8^. Injection experiments metadata for both macaque and marmoset can be found through the links provided in the **Key resources table**.

Labeled cells are counted and attributed to “source” areas according the species’ parcelation atlas and their position with respect to granular layer 4 (*i.e*. infragranular for layer 5 and 6; supragranular for layers 2 and 3). From this data extraction are computed two basic connectivity metrics. The first one is the Fraction of Labeled Neurons (FLN^75^), defined as the number of labeled cells in a given area divided by the total number of labeled cells for the injection (*i.e*. in all cortical areas, excluding labeling in the injected area; **Figure S1B**), thus giving the relative weight of the connection. By doing so, all injections are normalized to 1, thus allowing comparison between injections. The second metric is the Supragranular Labeled Neuron (SLN^76^), defined as the number of labeled cells in the supragranular compartment of an area divided by the total number of labeled cells for that area (*i.e*. both infra and supragranular; **Figure S1B**). This metric allows us to quantify the feedforward/feedbackness of a connection, with values close to 0 being identified as feedback (FB) and values close to 1 as feedforward (FF). These SLNs can be used to produce a species specific cortical hierarchy^15,51^ that has been shown to correlate with cortical gradients of several biological measurements, such as areal spine counts^31,53^ or the recently discovered neurotransmitter receptor gradients^32^. Over many injections, both metrics are usually displayed as matrices, with as many rows as there are areas in the relevant atlas (here macaque or marmoset), and as many columns as there is injections. Repeated injections are summed as if they where a single massive injection, which is consistent with standard practices when dealing with repeat injections, as in Markov et al. ^14,50^. In practice, what will be used throughout this paper is the square matrix (*i.e*. where source and target areas are the same) for each of those metrics, that is the edge-complete (*i.e*. all connections are known) graph of all commonly injected consensus areas.

#### Spine count gradient

The other anatomical constraint, besides connectivity, that is applied to the model is the gradient of spine count as investigated across several publications for both macaque^77–86^ and marmoset^82,87–89^. Spine counts, as captured by the total number of spine on the basal dendrites of layer 2/3 pyramidal cells, have been shown in macaques to vary across areas by more than tenfold from 600 in V1 more than 8000 in premotor areas. Further, Nimchinsky et al. ^90^ showed that approximately 90% of excitatory synapses on neocortical pyramidal cells are on dendritic spines, thus making a strong argument that a greater amount of spines results in a greater excitability. Elston ^91^ in turn suggested that spine count differences are likely to impact local summation of postsynaptic potentials. Because of this, spine counts can be used as a proxy to the self connectivity of a population.

Additionally, it is a non trivial discovery that these spine count values correlate positively with the SLN-based cortical hierarchy for both macaque and marmoset^8,31,53^, computed from retrograde tracer connectivity where the hierarchy is statistically inferred through a beta-binomial GLM applied to the SLN values^15,51^. This conception of hierarchy is itself rooted in the anatomical understanding of FB and FF^16,92,93^. The very fact of this correlation means that the hierarchy captures an important aspect of cortical areal differences, which can be used to complete the data where missing. All spine count values and their corresponding bibliographical references are given in **Table S1 and S2**, along with their hierarchical index.

#### Consensus mapping: Atlas

One of the key requirements of a formal comparative analysis of nervous systems is a step aimed at mapping the anatomical knowledge available for one species into registration with that available for the other. In the present study, this requirement translated into identifying homologous parcels of cerebral cortex in the marmoset and macaque, so the patterns of connectivity could be directly related within a same spatial reference frame. This process was, in practice, limited by issues related to the use of different nomenclatures by different laboratories, data availability (*e.g*. the areas explored with tracer injections may not be the same in the two species), and the likelihood of genuine biological differences. To circumvent such issues, we created a consensus map that describes the subdivision of the cortex of the two species in a manner that results in identification of the same number of homologous parcels (areas, or groups of areas) for which the results of tracer injections were available (**Figure 1, Table S3**).

This consensus mapping provides a way to directly relate the cortical areas proposed by Paxinos et al. ^94^ in the marmoset, which recognizes 116 histologically distinct areas, and those proposed by Markov et al. ^14^ in the macaque, which recognizes 91 areas. These atlases were chosen because they were used as the basis for describing the most comprehensive cellularlevel connectomic datasets available^8,9,14,15,60^. Alignment of the two parcelations was guided by several criteria, which included location in the brain and relative to other areas, histological structure, and, where available, functional data and pattern of connectivity.

In the simplest case, single areas could be directly identified in the two species using available histological and functional criteria. For example, the majority of the areas of the visual and premotor cortex could be identified with a high degree of certainty based on previous anatomical and functional studies^95,96^. In other cases, one area in one species was related to 2 or more areas in the other. For example, the marmoset caudal somatosensory cortex is not clearly differentiated into areas 1 and 2, as in the macaque, and appears instead the single area A1-2. Conversely, whereas marmoset area 3 is subdivided into A3a and A3b, the macaque atlas only provides a single subdivision called area 3. More complex mappings were required, leading to larger parcels. For example, in the macaque medial temporal lobe, area TH/TF was equated to five areas in the marmoset cortex: TF, TFO, TH, TL and TLO. At the end of this process, we are left with an consensus atlas of 75 areas. Let it here be noted that the subiculum (“SUBI” in **Figure 1**), entryway to the hippocampus proper and part of the hippocampal formation, is not part of the marmoset original parcelation, and therefore absent from the consensus one as well, effectively bringing down the consensus to 74 areas, since keeping the subiculum in the macaque is non consensual. Nonetheless, it was kept for completeness sake, and is not involved in the simulations reported here. The complete table of consensus equivalence across the two species is available in **Table S3**.

#### Consensus mapping: Computing connectivity

The consensus atlas now set, the connectivity of the two independent dataset must be converted to this new atlas of 74 common consensus areas, before they can be fully compared and used in a large-scale model. To do so, FLN, SLN and spine count values have to be combined so as to fit the new parcelation. As the consensus is essentially merging areas and area subdivisions together in either species to converge on homolog parcelation schemes, the connectivities of both species should reflect this by summing together source areas at the level of neuron counts. In other words, all the relevant connectivity transformations happen in the count matrices, created from the downloadable material for both species (see Method section **Anatomical connectivity data**). It must be stressed that to do so, one needs separated neuron count matrices for infragranular and supragranular numbers for both species, and that any operation needs to happen in the same fashion in both infra and supragranular matrices for each species.

For a given injection, two source areas (*i.e*. two rows in the count matrix) that need to be merged will see their respective counts of neurons summed together as a single area. Additionally, two target areas (*i.e*. two columns in the count matrix) will also be summed together, as if it were a large unique injection. The decision to sum, as opposed to average them together, follows the same logic as with the repeat injections explained above^14,50^. It stems from the fact that computing FLNs and SLNs are linear transformations that both involve dividing by the cumulated sum across injections. Additionally, averages tend to create non integer quantities for counts, where summing does not.

Once the new count matrices are ready and conform to the consensus parcelation, the macaque original matrices of 40×91 have become 35×74 matrices under consensus. Similarly, the marmoset ones have changed from 55×116 to 45×74. Complete consensus FLN and SLN matrices can be examined in **Figure S2 and S3** and full consensus data is available in **Table S5**. Of the 35 and 40 consensus injected areas for the macaque and the marmoset respectively, 29 are commonly injected in both species. Common consensus square *FLN* and *SLN* matrices of size 29×29 (**Figure 2 and S4**) are then extracted and, in the case of *FLN*, columns are renormalized to sum to 1, so that injections are directly compatible to one another. This is a standard practice in meso-connectomics, given the variation of injected volumes tracer from one injection to the next, and the sum total of neurons being a excellent proxy for injection volume and uptake zone size^75^.

#### Consensus mapping: Spine Count gradient

Merging spine counts to follow the consensus parcelation leads to two potential cases: either merging 2 areas (or more) that all have spine counts attributed to them prior to consensus, or merging 1 area that has a spine count with another area that does not. Through sheer happenstance, the first case never did present itself in either species, due in part to the fact that spine counts have been measured in the context of broader areas (*i.e*. area 46 or TE, which are larger than the currently used parcelation), and in part because far from all pre-consensus areas have been tested for spine counts. When the second case presented itself, the most parsimonious position was to assume homogeneity, and therefore attribute the spine count value of the first area to the resulting consensus area. This more parsimonious approach is further validated because spine counts are computed for an average neuron in each area.

In the case of consensus areas for which no spine count data was available prior to consensus, then the SLN-based hierarchy was used, itself computed from the consensus SLN matrices, as explained in Method section **Spine count gradient** and **Computational model: Spine count gradient as synaptic strength**, following the methods from Vezoli et al. ^51^ and Markov et al. ^15^.

#### Consensus mapping: 11×11 matrices

In order to study the particularities of the WM system proper, we expended the 2-compartment model of Murray et al. ^40^, which modeled PPC and DLPFC with each with a single Wong-&-Wang, into an 11 area model, where PPC and DLPFC are now 6 and 5 areas, respectively. Doing so therefore encompasses all available consensus data regarding the parietofrontal network, known to be the substrate of WM^27^. These 11×11 matrices can be seen in **Figure 4B** and consist of 6 PPC and 5 DLPFC consensus areas. The corresponding spine count values were taken from the consensus spine count gradient described above in Method section **Consensus mapping: Spine Count gradient**.

#### Computational model: local circuit

Conceptually, the system built here should be thought about as a network of Wong-&-Wang models^31,68^, each corresponding to a particular cortical area, and whose connectivity to other areas (*i.e*. other Wong-&-Wangs) are structured and constrained by the retrograde tracer connectivities described above (**Figure 3A**).

The local system, which belongs to the category of mean-field models, possesses in our case three explicit variables *r*_*A*_, *r*_*B*_ and *r*_*C*_, describing the temporal evolution of the firing rates of three neural populations: two excitatory (populations A and B) and one inhibitory (population C). The equation of the firing rate for each population is as follows:

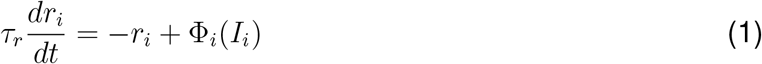

Here, and in all future equations, all parameter values are taken from Mejias & Wang^31^, except *G* and *ρ*. The index *i* can be A, B or C, depending on the population. *τ*_*r*_ = 2 ms and Φ_*i*_(*I*_*i*_) is a transfer function that converts the total current *I* of a population into a firing rate, according to the following equations^97^:

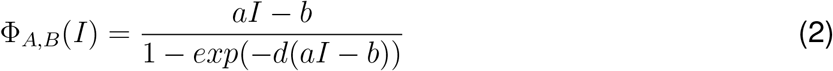

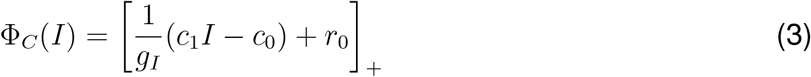

with *a* = 135 Hz/nA, *b* = 54 Hz, and coefficient *d* = 0.308 for equation (2); and *g*_*I*_ = 4, *c*_1_ = 615 Hz/nA, *c*_0_ = 177 Hz and *r*_0_ = 5.5 Hz for equation (3). The index “+” in [*expr*]_+_ indicates a rectification where all negative values are set to 0.

Given that Φ_*i*_ evolves as a function of *I*_*i*_, let us define next the current equations for each population:

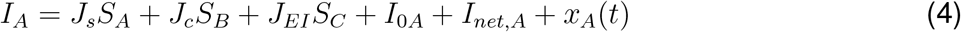

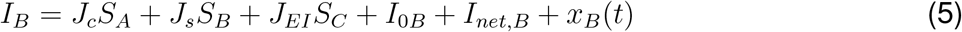

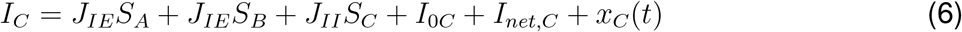

Starting with the fixed parameters for this set of equations, the cross-coupling parameter *J*_*c*_ has a value of 0.0107 nA and encapsulates how the two excitatory populations A and B crossactivate each other. *J*_*EI*_ is the coupling parameter regulating the input from inhibitory population C into excitatory populations A and B. Being inhibitory, the term *J*_*EI*_*S*_*C*_ must be negative, so *J*_*EI*_ is set at −0.31 nA. *J*_*II*_, population C inhibitory input onto itself, is also negative at −0.12 nA. Background current parameters *I*_0*A*_ and *I*_0*B*_ are identical and set at 0.3294 nA, *I*_0*C*_ is set at 0.26 nA.

Although fixed, the self-coupling parameter *J*_*s*_ and the excitatory-to-inhibitory coupling parameter *J*_*IE*_ are different for each area depending on the spine count gradient (see Method section **Spine count gradient**) and will be explained in a dedicated section down below (Method section **Computational model: Spine count gradient as synaptic strength**). Similarly, the *I*_*net,i*_ terms for each populations, which encapsulates the role of the inter-areal anatomical connectivity (Method section **Anatomical connectivity data**), requires its own dedicated explanations further down (Method section **Computational model: Inter-areal connectivity**).

The variables collectively designated as *S*_*i*_ are described by the conductance equations below, and truly govern the local system:

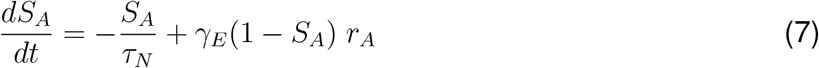

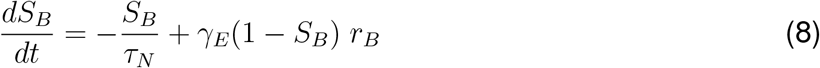

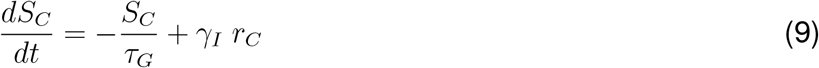

In these, *S*_*i*_ are the conductances for each population; the *τ*_*N*_ (NMDA) and *τ*_*G*_ (GABA) time constants are set at 60 and 5 ms respectively; and *γ*_*E*_ and *γ*_*I*_, set at 1.282 and 2, are dimensionless saturation factors for the excitatory and inhibitory populations, respectively again. Finally, *r*_*i*_ is the firing rate of each population from equation (1).

The last term in the current equations (4-6), *x*_*i*_(*t*), signifies an Ornstein-Uhlenbeck process, which introduces some level of stochasticity, and can be construed as random noise intrinsic to the system. It is defined be the following equation:

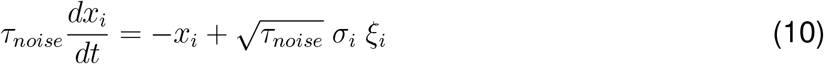

where *τ*_*noise*_ is the time constant for the process set at 2 ms, *σ*_*i*_ is the noise strength set at 0.005 nA for populations A and B, at 0 for population C. The last term *ξ*_*i*_ is the Gaussian white noise, on which *σ*_*i*_ is applied.

Finally, still in equations (4-6), an implicit term *µ* can be added to introduce sensory stimulations into the system, in which case *µ* = 0.3 nA for a duration of 500 ms. **Figure 4D** shows the schematic progression of the task modeled in this study, with the cue and the distractor stages being where *µ* is non zero for one of the populations.

#### Computational model: Spine count gradient as synaptic strength

What makes large-scale models increasingly important as neuroscience progresses is their ability to incorporate differences between cortical areas, where they would otherwise be construed — and modeled — as rigorously identical units. Here, we use a brain wide gradient of spine count to uniquely constrain our system, whereby each area is characterized by a spine count value, as exposed in Method section **Spine count gradient**, which is used to compute an area-specific self-coupling parameter *J*_*s*_ through the following equation:

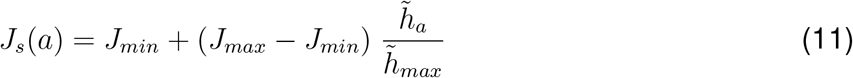

where *a* is a consensus area; *J*_*min*_ is the minimal synaptic strength set at 0.21 nA; and *J*_*max*_ is the maximal value synaptic strength can take, set at 0.42 nA, effectively bounding *J*_*s*_ between those to values. 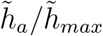 is meant to represent the normalized spine count gradient, completed by SLN-based hierarchical estimates 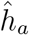 where necessary (see **Table S1**). This effectively produces a gradient of *J*_*s*_ values, spanning from *J*_*min*_ to *J*_*max*_. *J*_*max*_ being inferior to bifurcation threshold *J* ^*∗*^ ensures that no single area taken alone is able to achieve multistability (**Figure 4C**).

From *J*_*s*_ is computed *J*_*IE*_ with the subsequent equation:

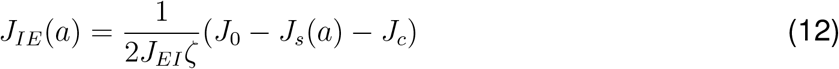

where *a* is again an area; *J*_*c*_ and *J*_*EI*_ are the same parameters as in equations (4-6), set at 0.0107 and −0.31 nA respectively. Parameter *ζ* is fixed on parameters previously spelled out:

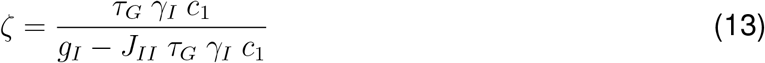

with *τ*_*G*_ and *γ*_*I*_ from equation (9); *c*_1_ and *g*_*I*_ from equation (3); and *J*_*II*_ from equation (6). The explanation for the precise definition of *ζ* can be found in the methods of Mejias & Wang^31^. Briefly, the baseline activity level of each local Wong-&-Wang circuit is by default dependent on *J*_*s*_, and areas with high values could exhibit spontaneous persistent activity. They could in principle even go beyond physiologically realistic limits if left unbounded. We control this by imposing on the system that all areas have the same baseline, regardless of their specific *J*_*s*_ value. We allowed three simplifying assumptions: The system is symmetrical with respect to populations A and B (at baseline, the only input the background current, and *I*_0*A*_ = *I*_0*B*_), noise is negligible, and population C has significantly faster dynamics, as they are mediated by GABA. After simplification, equation (12) and (13) are deduced, with *J*_0_ being set at 0.2112 nA. The linear relationship described in equations (11) and (12) ensures that the baseline activity is the same for all areas in the network. Note that deviations from this linear relationship would simply lead to different areas having slightly different spontaneous activity levels, but it does not substantially affect our main results.

Because *J*_*IE*_ needs to always be non negative, equation (12) imposes a lower bound to the values *J*_*s*_ can take in equation (11) at 0.205 nA. On the other side of its range, a *J*_*s*_ that would be higher than the bifurcation point of the system (0.4655 nA for an isolated Wong-&-Wang parametrized as described so far) would grant multistability to the system. Therefore, setting *J*_*max*_ below that bifurcation threshold ensures that all areas are monostable when taken in isolation. In other worlds, each individual area cannot on its own display sustained activity, even after a stimulation, and any persistent non baseline firing rate is strictly an effect multi-areal system as a whole. Here, as described above, setting *J*_*min*_ and *J*_*max*_ at 0.21 and 0.42 nA respectively ensures that our system displays distributed WM as a non linear effect of associating monostable areas together.

#### Computational model: Inter-areal connectivity

Going back to equations (4-6), the term *I*_*net,i*_ still requires explanations. This term designates the current that, for each population in each area, comes from the rest of the network outside the area in question. Said differently, it is the global input current to each area, and is defined as such:

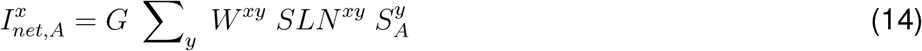

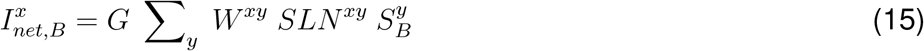

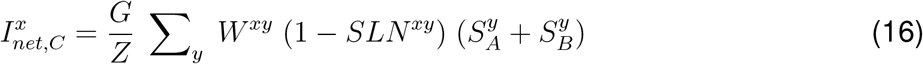

In these, indices *x* and *y* respectively represent rows and columns of matrices *W*^*xy*^ and *SLN*^*xy*^, where each column is a target area (*i.e*. injection site) and each row is a source area. Additionally, *y* the *y*^*th*^ element in column vector 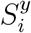. The summation over *y* applies over the expression 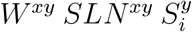 (or the population C equivalent), which is the pairwise (*i.e*. Hadamard) product of matrices *W, SLN* and column vector *S*_*i*_. The output of said summation is the row vector 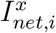, where *x* is the *x*^*th*^ element in the vector.

*G* stands for the global coupling parameter, which tunes up or down the importance of the input from the network to a given area. *G* has to scale down as the number of areas scales up, or the sum that is *I*_*net*_ will gradually take over the current equation (4-6) and the dynamics will blow up exponentially. The exact value of *G* is different for the two species (see Results: Standard Behavior) and are set at 0.98 and 0.85 (0.4 and 0.42 in the 2×2 case; 0.51 and 0.33 in the 29×29 case) for macaque and marmoset respectively, for all simulations. *Z* is a scalar that controls the balance between excitatory and inhibitory inter-areal connections. *Z* = 1 would mean that long-range excitatory and inhibitory projections are of equal strength. In our particular case, we needed to enforce the following condition: if excitatory populations A and B are both active at the same time and at the same level in a given area, then their net effect on other areas in the network should be 0. Given this imperative, parameter *Z* can be deduced as follows:

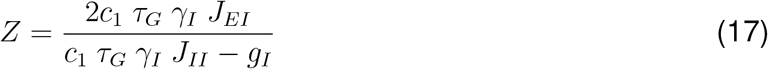

with *c*_1_ and *g*_*I*_ from equation (3); *τ*_*G*_ and *γ*_*I*_ from equation (9); *J*_*EI*_ and *J*_*II*_ from equations (4-6).

Back to equations (14 to 16), *W*^*xy*^ is the weight matrix, which we get to through the following sequence of transformations: First we define the matrix *U* as the non linear rescaling of matrix *FLN* (see Method section **Anatomical connectivity data**):

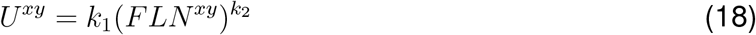

where *k*_1_ and *k*_2_ are rescaling factors of value 1.2 and 0.3 respectively. The rescaling described by equation (18) is done so that the range of FLN values, which spans 5 to 6 orders of magnitude, is better suited to a firing rate model. The same qualitative behavior can be obtained with other *k*_1_ and *k*_2_ values, or even other rescaling functions, so long as a standard working regime is achieved. The values were kept from Mejias & Wang^31^ for consistency.

We then renormalize *U* by column to obtain *V*:

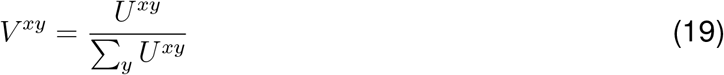

Importantly, this renormalization step, named as such to differentiate it from the original normalization that produces FLNs, is to be performed only once, on the edge-complete 29×29 matrix. The 11×11 connectivity matrix of parietal and prefrontal areas is not renormalized again, as it is considered a subpart of a larger network, not its own independent system.

Finally, to obtain our true weight matrix *W*, the spine count gradient is introduced to the inter-areal connectivity so that global and local circuitry be of the same trend, like so:

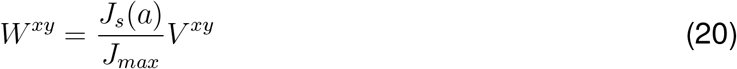

Going back once more to the *I*_*net,i*_ equations (14-16), the *SLN* term indicates the square matrix of SLN values, already described in **Anatomical connectivity data**. As SLN values evolves between 0 and 1, with 0 meaning FB and 1 as FF, pairwise multiplication of the *SLN* and the *W* matrices gradually filters out FBs as a function of the SLN value. If it is a reasonable assumptions that FFs should be excitatory, as it facilitates signal propagation to the entire system, having FB do the same would lead to excitatory reverberation and all areas being indiscriminately activated as well as having similar firing rates during WM delay, as shown by Mejias & Wang^31^. In the same paper, the authors introduce the notion of Counterstream Inhibitory Bias (CIB), by which the weight matrix *W* is, for the inhibitory population C of each area, not pairwise multiplied by *SLN* but by (1 −*SLN*), thus filtering out FF connections, not FB ones. By doing so, the model effectively takes on the assumption that FBs connect to inhibitory neurons which, if it is very much consistent with the known FB modulatory effects^98–100^, the existence of such FB-to-inhibitory-cell connections still remains to be proven at the cellular level. That being said, applying the CIB to a distributed WM model does yield a pattern of sustained activity during WM that is remarkably close to known electrophysiological data^31^, thus making the CIB a fascinating as well as credible prediction to be tested by experimentalists.

Finally, all parameters of the model are listed in **Table S4**.

## Quantification and statistical analysis

### Connectome statistics and SLN-based hierarchy

All Statistics were performed in R. Macaque vs marmoset FLN correlations (**Figure 3**) where done using the linear model function of the *stats* package, which is part of the base sets of packages in R. The SLN-based hierarchy of each species was computed following the methods developed in Markov et al, 2014^15,51^. In short, we suppose that we can assign hierarchical levels, *h*_*i*_ and *h*_*j*_, to all area pairs *i* and *j*, based on a the SLN measurements taken from injection in area *a*. Then, we assume that:

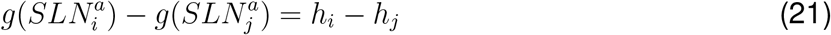

where *g* applies a logit or probit transformation to SLN values, from an injection into area *a* that receives projections from areas *i* and *j*. This suggests a formalism similar to a GLM with a binomial family. The SLN is taken as a binomial variable (neurons are found in the upper or lower cortical layers) and the sum of neurons in both compartments is used a weight. The problem can be reframed as follows:

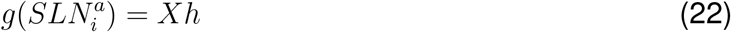

where *X* is the incidence matrix of the cortical graph, and *h* the vector of hierarchical values, the product of which maps the differences in hierarchical value between two areas with the differences between the transformed SLN.

A negative beta-binomial model (with a probit link function) was used, given that the distribution of the response variable, here the expected SLN values, has a greater variability than the usual binomial distribution and is better described by a beta distribution (aods3 linear model from the *aods3* R package^101^). For present purposes, the model can be reparameterized to include a dispersion parameter that better models the overdispersion typically observed in neural counts^15^. Once the statistical model is specified, the coefficients are estimated by maximum likelihood. Note that because numbers of neurons are used in the model and not just the SLN proportions, this method generates a weighted hierarchy.

## SUPPLEMENTARY MATERIAL

### Supplementary figures

**Figure S1:**
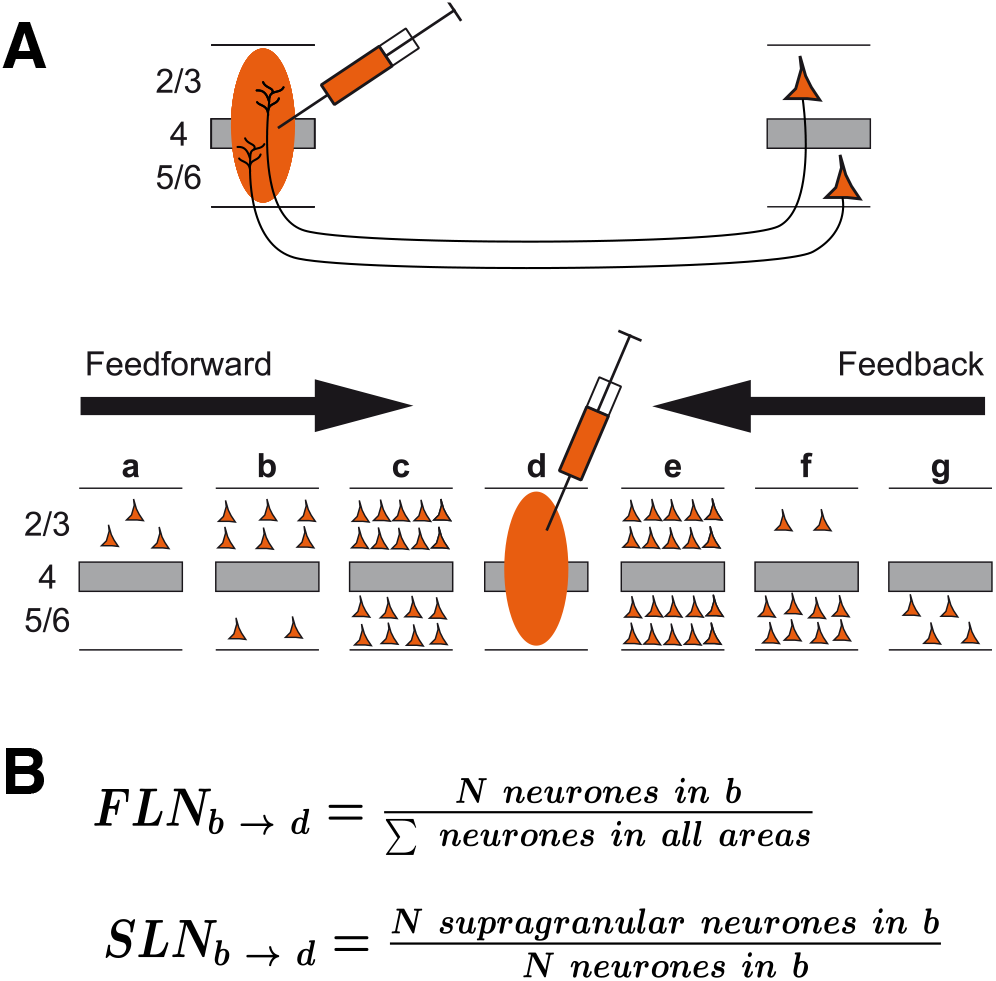
Retrograde tract tracing and connectivity computations. (A) In retrograde tract tracing, the tracer is captured by axon terminals present in the uptake zone (orange oval) and actively transported back to the soma of the cells. FF connections (*i.e*. from lower hierarchical levels) originates predominantly in the supragranular layers, a dominance that becomes gradually more apparent as hierarchical distance increases. The converse is symmetrically true for FBs and infragranular layers. (B) FLN and SLN equations to compute both metrics for each existing connection.

**Figure S2:**
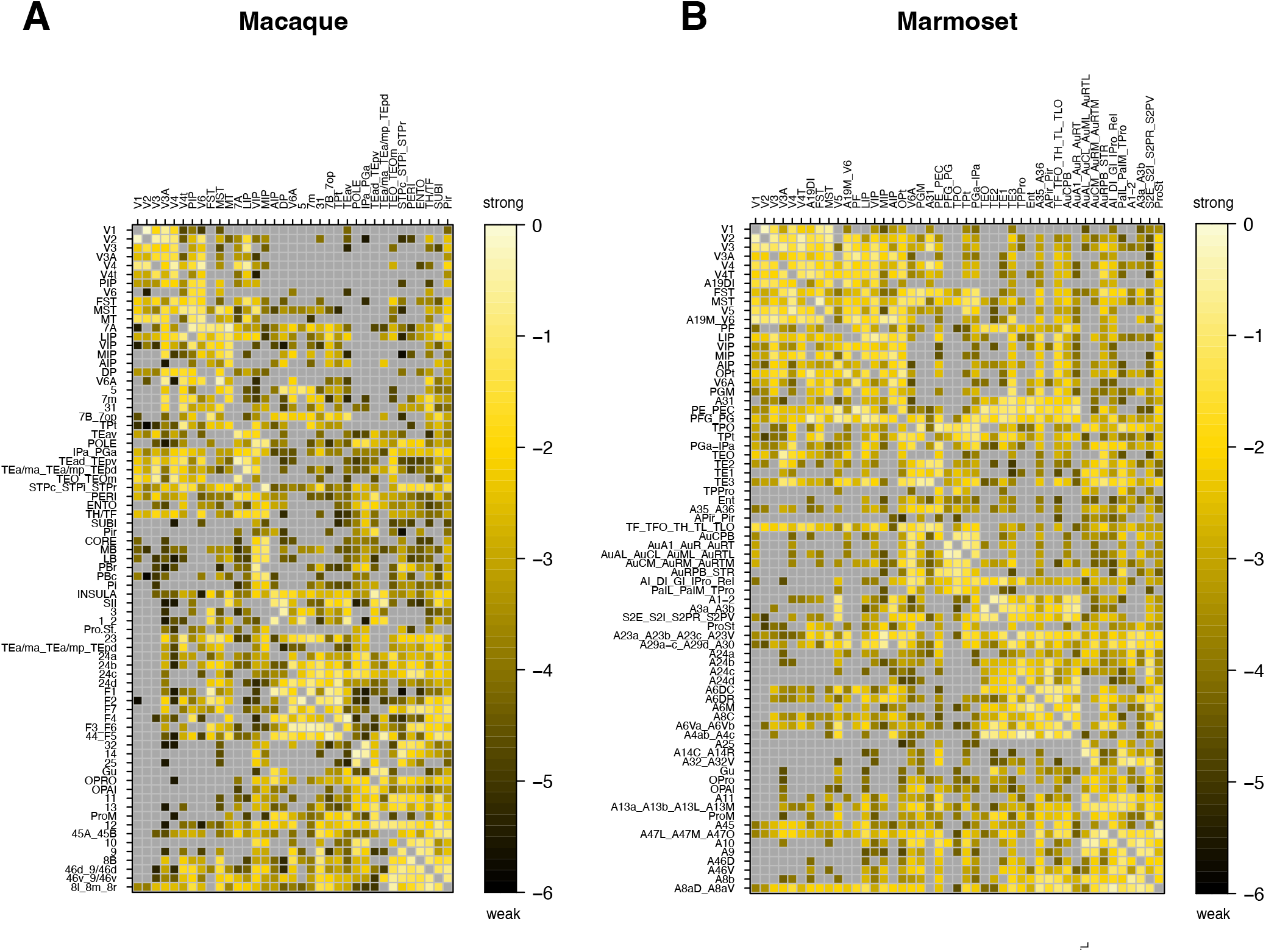
Full consensus FLN connectivity matrices. The consensus mapping reduces both species’ atlases to a common parcellated of 74 areas, plus the subiculum (“SUBI”) in the macaque. Reading convention as in **Figure 2**. (A) After consensus, the macaque goes from a 40×91 matrix to a 35×75 matrices. (B) Similarly, the marmoset goes from 55×116 to 45×74.

**Figure S3:**
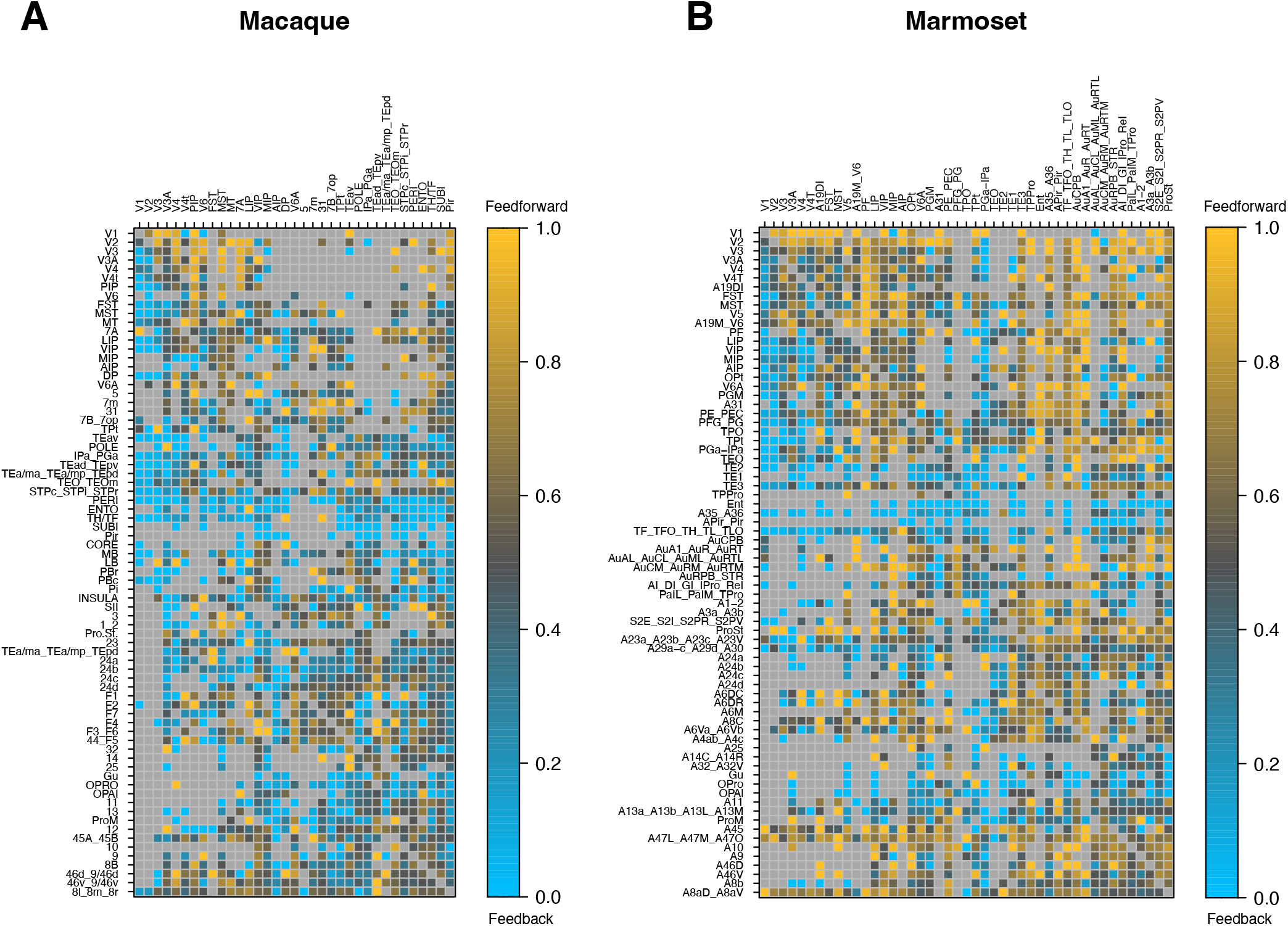
Full consensus SLN connectivity matrices. Same dimensions as in **Figure S2**. SLNs vary between 0 (FB; blue hues) and 1 (FF; yellow gold hues). (A) Macaque. (B) Marmoset.

**Figure S4:**
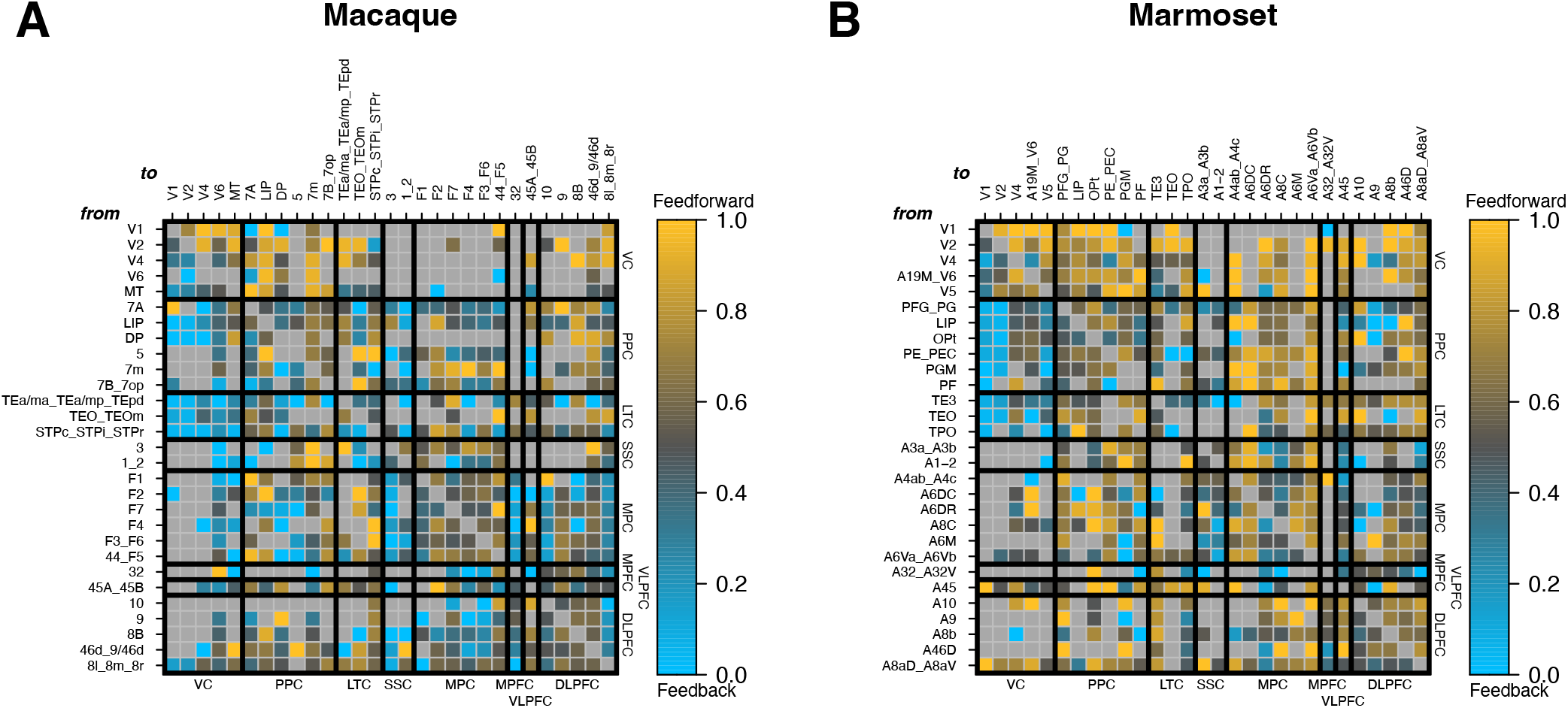
29×29 common consensus SLN matrices for both species. All connections are here one-to-one comparable across species. Reading convention as in **Figure S3** (A) Macaque. (B) Marmoset.

**Figure S5:**
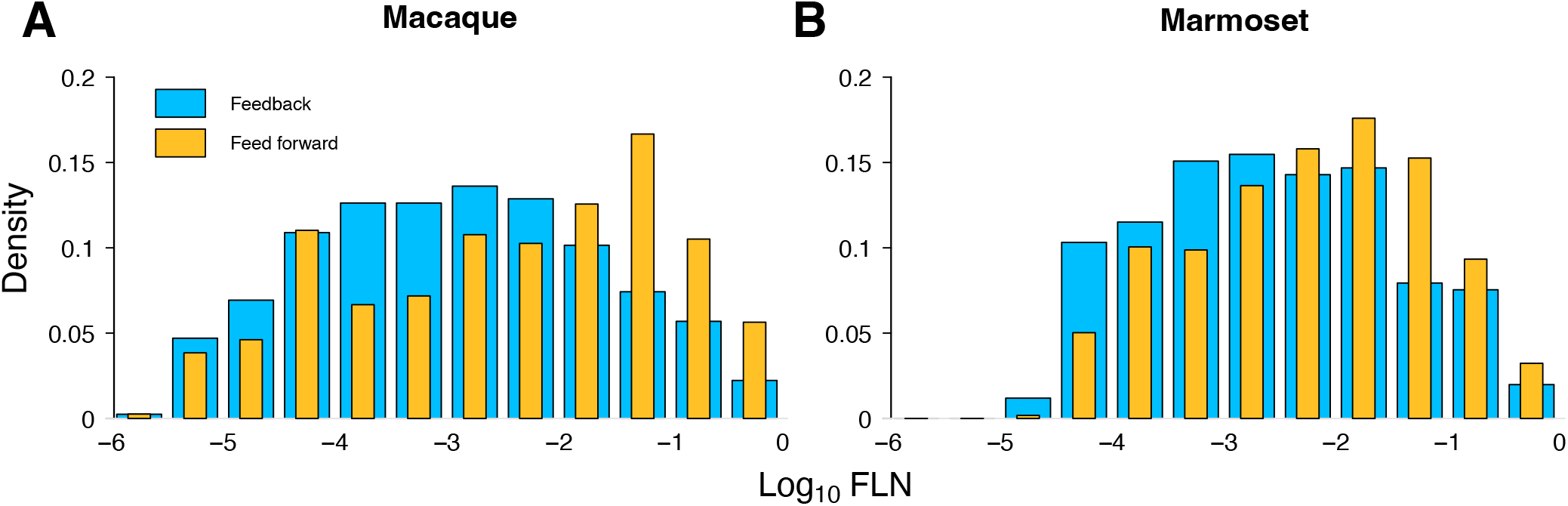
Feedback (FB) and feedforward (FW) distribution as a function of *log*_10_ FLN. Blue, feedback ; yellow, feedforward. (A) Macaque. (B) Marmoset.

**Figure S6:**
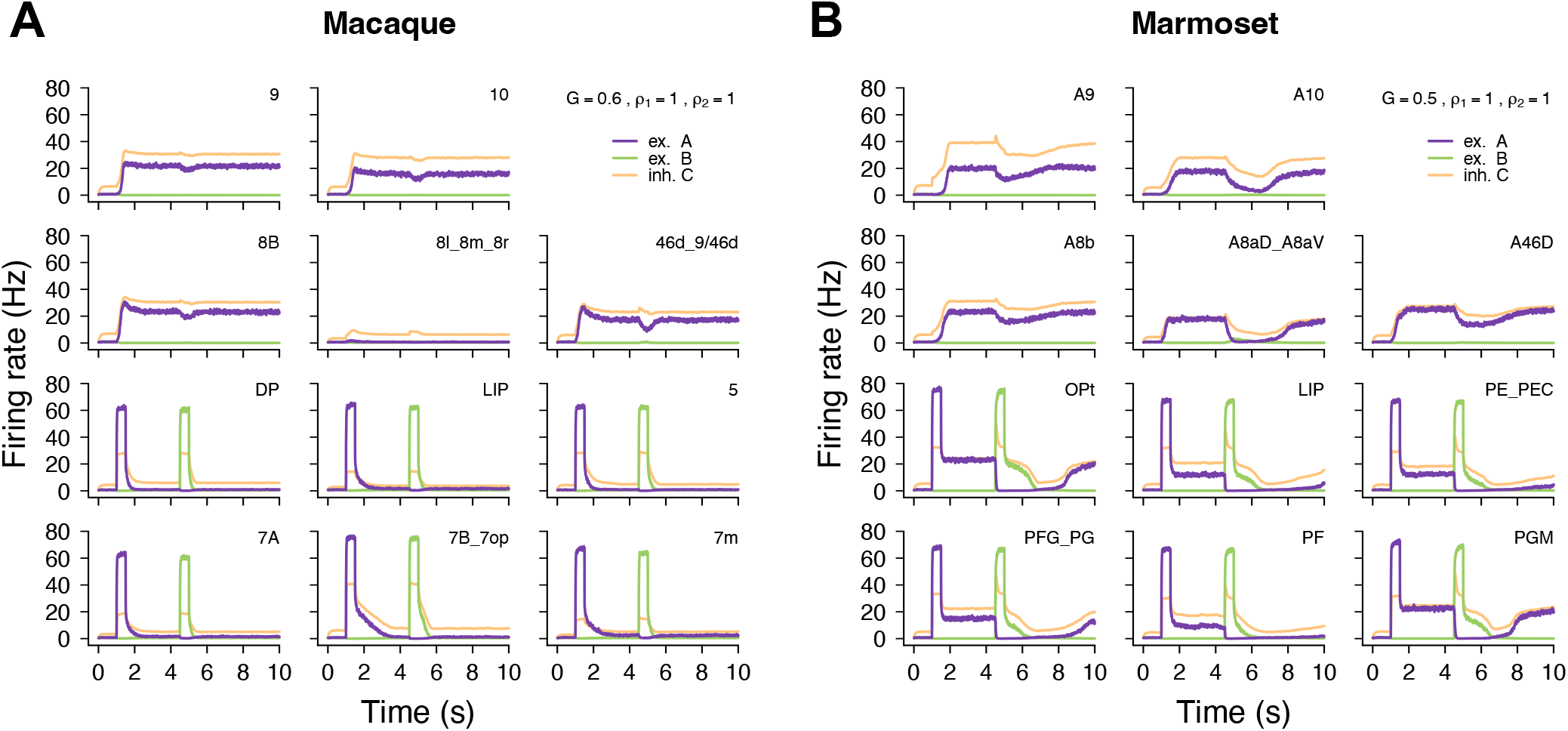
Alternative parametrization of global coupling parameter *G*. (A) Full example of firing rate as a function of time for the macaque’s partially persistent regime at *G* = 0.6 from **Figure 5B**. (B) Full example behavior for the marmoset’s small window of resilient regime at *G* = 0.5 from **Figure 5C**. Reading conventions as in **Figure 5B and C**.

**Figure S7:**
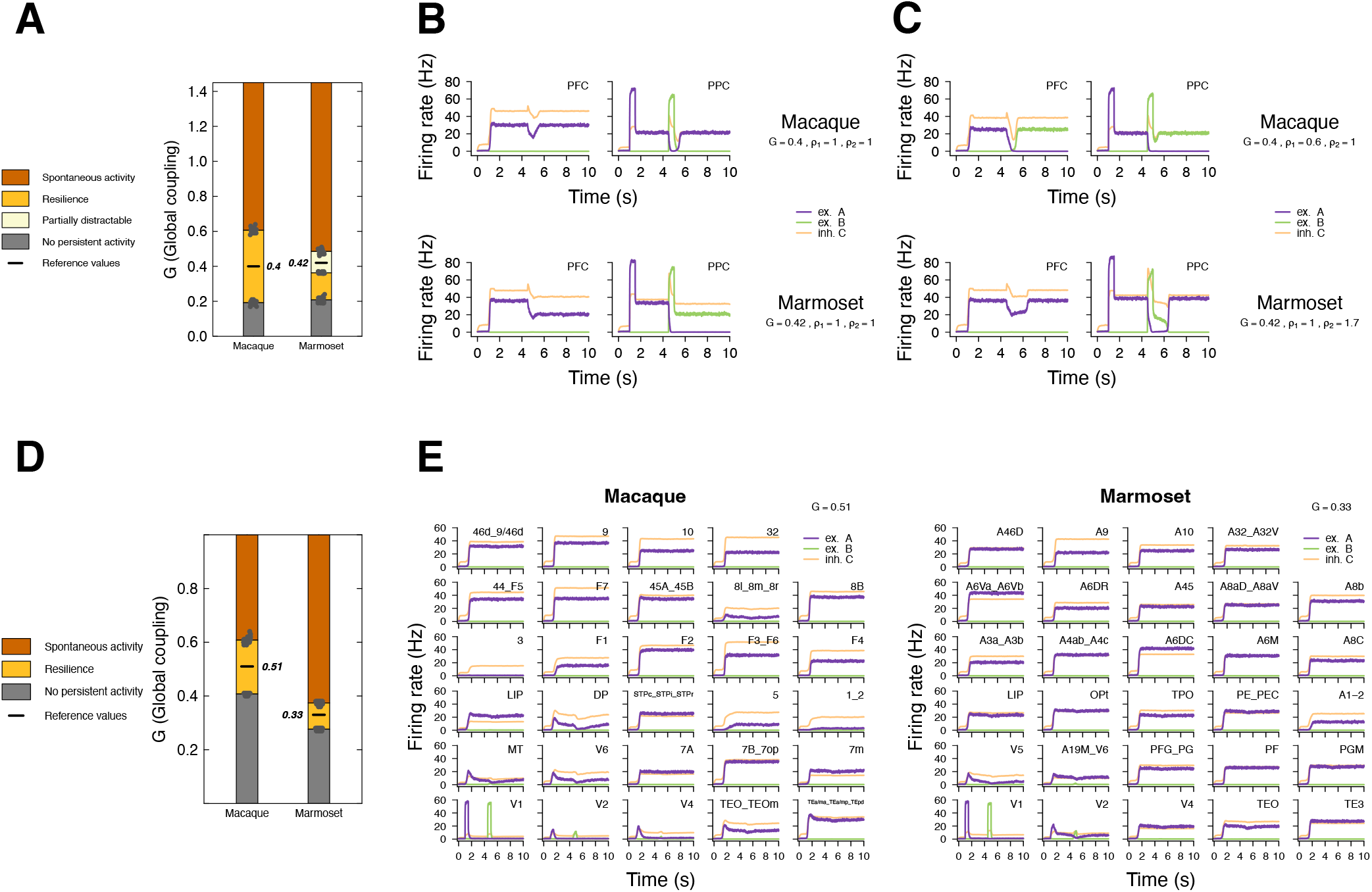
2×2 PFC-PPC and 29×29 area model of the distributed frontoparietal working memory model. (A) Stacked histogram showing different regimes of activity as a function of *G*, for both species, in the 2×2 case, computed as in **Figure 5A**, over 20 different random seeds. (B) 2×2 PFC-PPC standard behavior with global coupling parameter *G* fixated at 0.4 for the macaque and 0.42 for the marmoset. The marmoset (lower panel) shows partial sensitivity to distraction. (C) A distractible macaque can be achieved by adjusting *ρ*_1_ to 0.6, as in the 11×11 case (upper panel). A resilient marmoset arises by adjusting *ρ*_2_ to 1.7 (lower panel), consistent with parameter space in **Figure 7A**. (D) Stacked histogram showing different regimes of activity as a function of *G*, for both species, in the 29×29 case, computed as in **Figure 5A**, over 20 different random seeds. (E) 29×29 standard behavior with global coupling parameter *G* fixated at 0.51 for the macaque and 0.33 for the marmoset. Both species are resilient. Reading conventions for (A) and (D) as in **Figure 5A**; reading conventions for (B), (C) and (E) as in **Figure 5B and C**.

**Figure S8:**
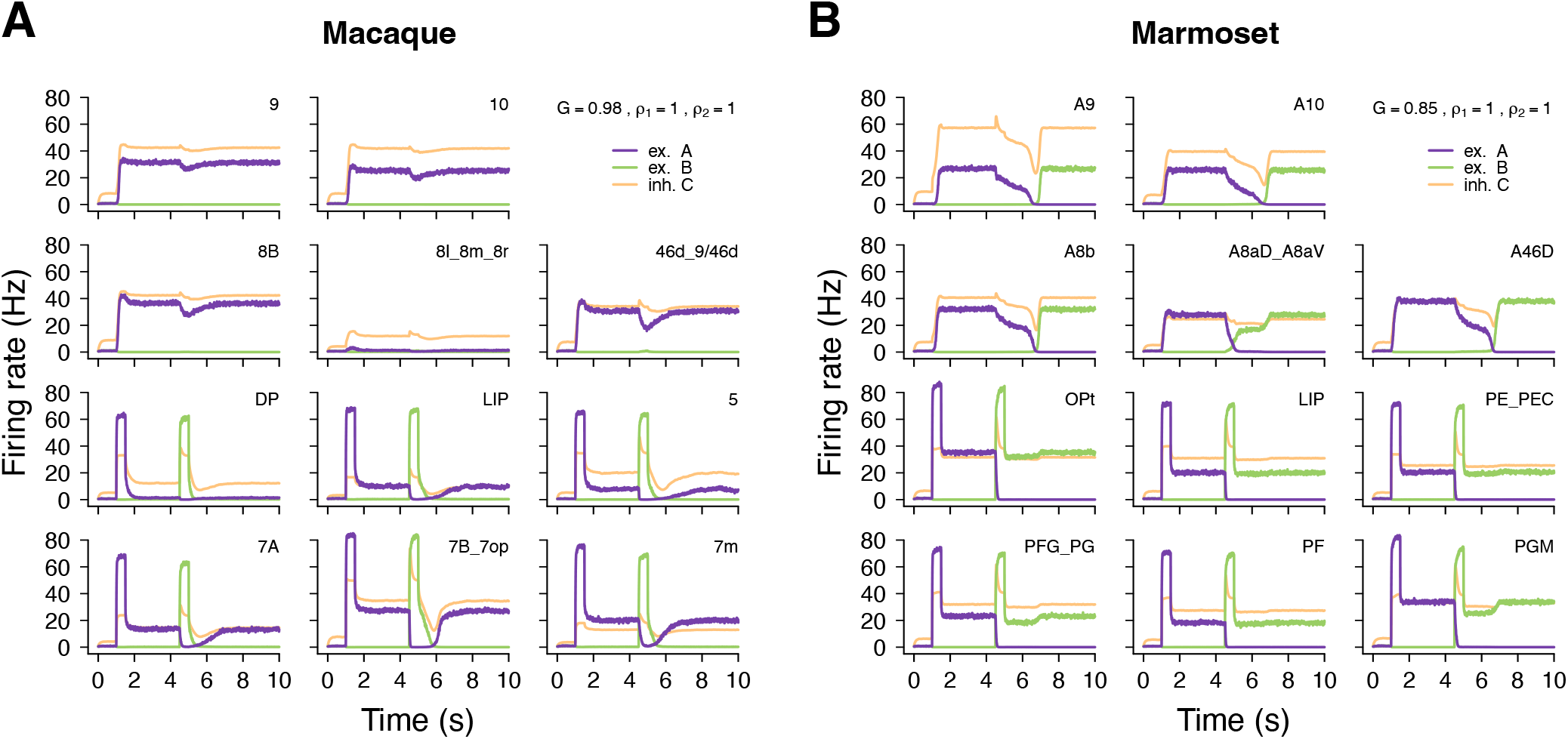
Model’s behavior after swapping species specific connections. (A) The macaque connections (FLN and SLN) that do not exist in the marmoset were removed and replaced by the marmoset connections that do not exist in the macaque. Connections that exist in both were left unchanged. (B) Same procedure for the marmoset, performed in the opposite direction. *G* parameters unchanged. Reading conventions as in **Figure 5B and C**.

**Figure S9:**
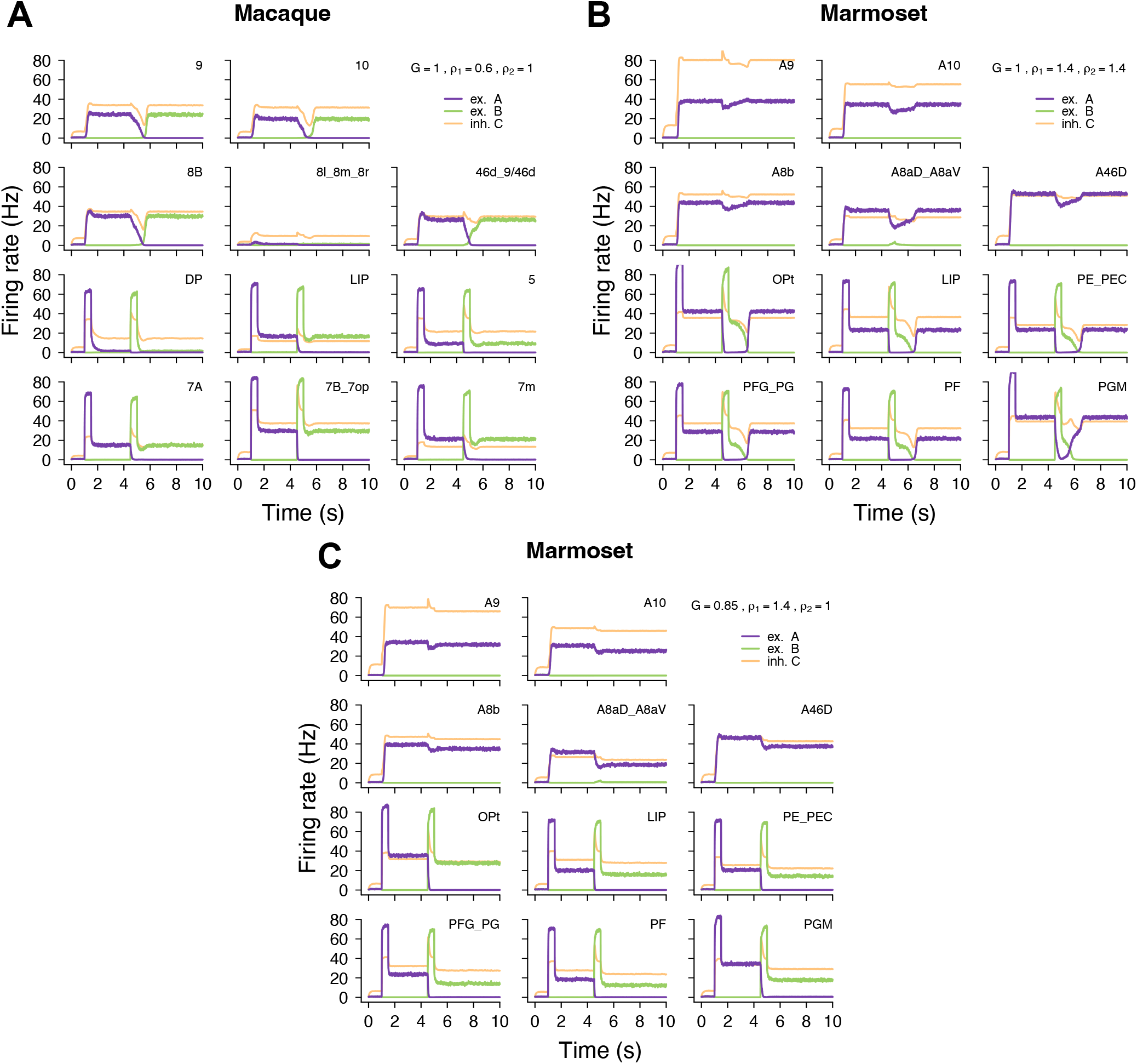
*ρ*_1_− *ρ*_2_ Parameter space examples of resilience, distractibility and mixed behavior. (A) Full example behavior for the macaque’s distractible regime at *G* = 0.98, *ρ*_1_ = 0.6, *ρ*_2_ = 1. (B) Full example behavior for the marmoset’s resilient regime at *G* = 0.85, *ρ*_1_ = 1.4, *ρ*_2_ = 1.4. (C) Full example behavior for the marmoset’s mixed regime at *G* = 0.85, *ρ*_1_ = 1.4, *ρ*_2_ = 1.

**Figure S10:**
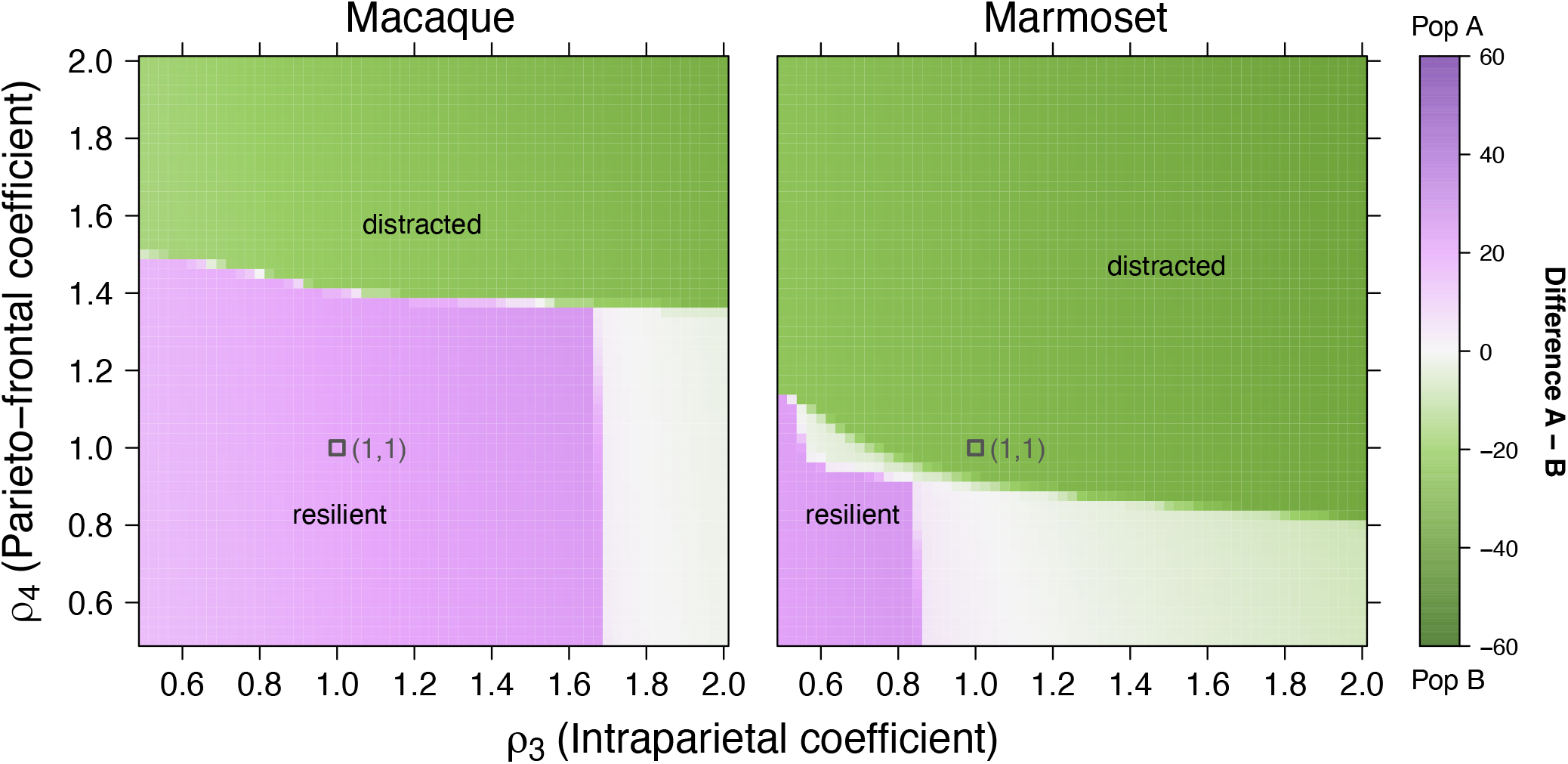
*ρ*_3_ − *ρ*_4_ Parameter space analysis of resilience and distractibility. Reading conventions as in **Figure 7B**). Each square is the average of simulations performed over 10 different random seeds, by incremental steps of 0.025 in both directions. Black open squares indicate (1,1) coordinates in the plane.

### Supplementary tables

**Table S1:** Macaque spine count, hierarchical values and literary sources. 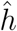 are the SLN based hierarchical position estimates. For V1, area 10, and consensus areas 46d 9/46d and TEa/ma TEa/mp TEpd, spine count values at the top are the mean of the values held between angular brackets just below. The last column indicates the name of the area for which the count was made in its original atlas conventions.

| Consensus area | $\hat{h}$ | Spine count | Source | Area name in source |
| --- | --- | --- | --- | --- |
| V1 | 0 | 643<br>(688, 597) | Elston et al. 1999<br>Elston and Rosa, 1997 | V1 |
| V2 | 0.132269 | 1201 | Elston and Rosa, 1997 | V2 |
| V4 | 0.374088 | 2429 | Elston and Rosa, 1998b | V4 |
| 3 | 0.377149 | 2987 | Elston and Rockland 2002 | 3b |
| V6 | 0.488866 |  |  |  |
| 1_2 | 0.552926 |  |  |  |
| DP | 0.601343 |  |  |  |
| MT | 0.613831 | 2077 | Elston and Rosa 1997 | MT |
| TEO_TEOm | 0.641515 | 4800 | Elston et al. 2010a | TEO |
| 5 | 0.673251 | 4689 | Elston and Rockland 2002 | 5 |
| 8l_8m_8r | 0.68387 | 3200 | Elston and Rosa, 1998a | FEF |
| F1 | 0.709641 | 4568 | Elston and Rockland 2002 | 4 |
| F4 | 0.735719 |  |  |  |
| F3_F6 | 0.792739 | 8238 | Elston and Rockland 2002 | 6 |
| 32 | 0.800469 |  |  |  |
| 10 | 0.825804 | 6488<br>(6887, 6090) | Elston et al. 2011a | 10 |
| 44_F5 | 0.843874 | 8238 | Elston and Rockland 2002 | 6 |
| 7A | 0.851916 | 2572 | Elston and Rosa 1997 | 7a |
| 45A_45B | 0.858629 |  |  |  |
| 46d_9/46d | 0.858915 | 6584<br>(6621, 6548) | Elston et al. 2011a | 46 |
| LIP | 0.869096 | 2316 | Elston and Rosa 1997 | LIPv |
| 7m | 0.904036 | 2294 | Elston 2001 | 7m |
| 7B_7op | 0.904156 | 6841 | Elston and Rockland 2002 | 7b |
| F7 | 0.92325 | 8238 | Elston and Rockland 2002 | 6 |
| 8B | 0.923584 |  |  |  |
| STPc_STPi_STPr | 0.924344 | 3585 | Elston 2001 | STPp |
| 9 | 0.947519 | 7637 | Elston et al. 2011a | 9d |
| F2 | 0.981861 | 8238 | Elston and Rockland 2002 | 6 |
| TEa/ma_TEa/mp_TEpD | 1 | 6730<br>(7260, 6200) | Elston et al. 1999 (TEad)<br>Elston et al. 2009 (TEpd) | TE |

**Table S2:** Marmoset spine count, hierarchical values and literary sources. Reading conventions as in **Table S1**.

| Consensus area | $\hat{h}$ | Spine count | Source | Area name in source |
| --- | --- | --- | --- | --- |
| V1 | 0 | 950 | Oga et al. 2013 | V1 |
| V2 | 0.180824 | 1240 | Elston et al. 1999 | V2 |
| V5 | 0.429611 | 2359 | Elston et al. 1999 | MT |
| A19M_V6 | 0.508004 | 1338 | Elston et al. 1999 | DM |
| A9 | 0.565519 | 4843 | Sasaki et al. 2015 | 8B/9 |
| A10 | 0.571427 | 3983 | Elston et al. 2001 | 8B/9 |
| TEO | 0.593298 | 3465 | Elston et al. 1999 | TEO |
| A1-2 | 0.606556 |  |  |  |
| V4 | 0.643048 | 2098 | Elston et al. 1999 | DL |
| A45 | 0.645245 |  |  |  |
| A8aD_A8aV | 0.691981 |  |  |  |
| PF | 0.692536 |  |  |  |
| A6DR | 0.693295 |  |  |  |
| LIP | 0.709286 |  |  |  |
| A3a_A3b | 0.723727 |  |  |  |
| A8C | 0.732859 |  |  |  |
| PE_PEC | 0.737931 |  |  |  |
| A46D | 0.747617 |  |  |  |
| TE3 | 0.748794 | 3100 | Oga et al. 2013 | TE3 |
| TPO | 0.761058 |  |  |  |
| PFG_PG | 0.768203 |  |  |  |
| PGM | 0.777864 |  |  |  |
| A6M | 0.791358 |  |  |  |
| OPt | 0.805353 |  |  |  |
| A8b | 0.809164 | 4843 | Sasaki et al. 2015 | 8B/9 |
| A4ab_A4c | 0.819596 |  |  |  |
| A32_A32V | 0.823476 |  |  |  |
| A6DC | 0.93911 |  |  |  |
| A6Va_A6Vb | 1 |  |  |  |

**Table S3:**
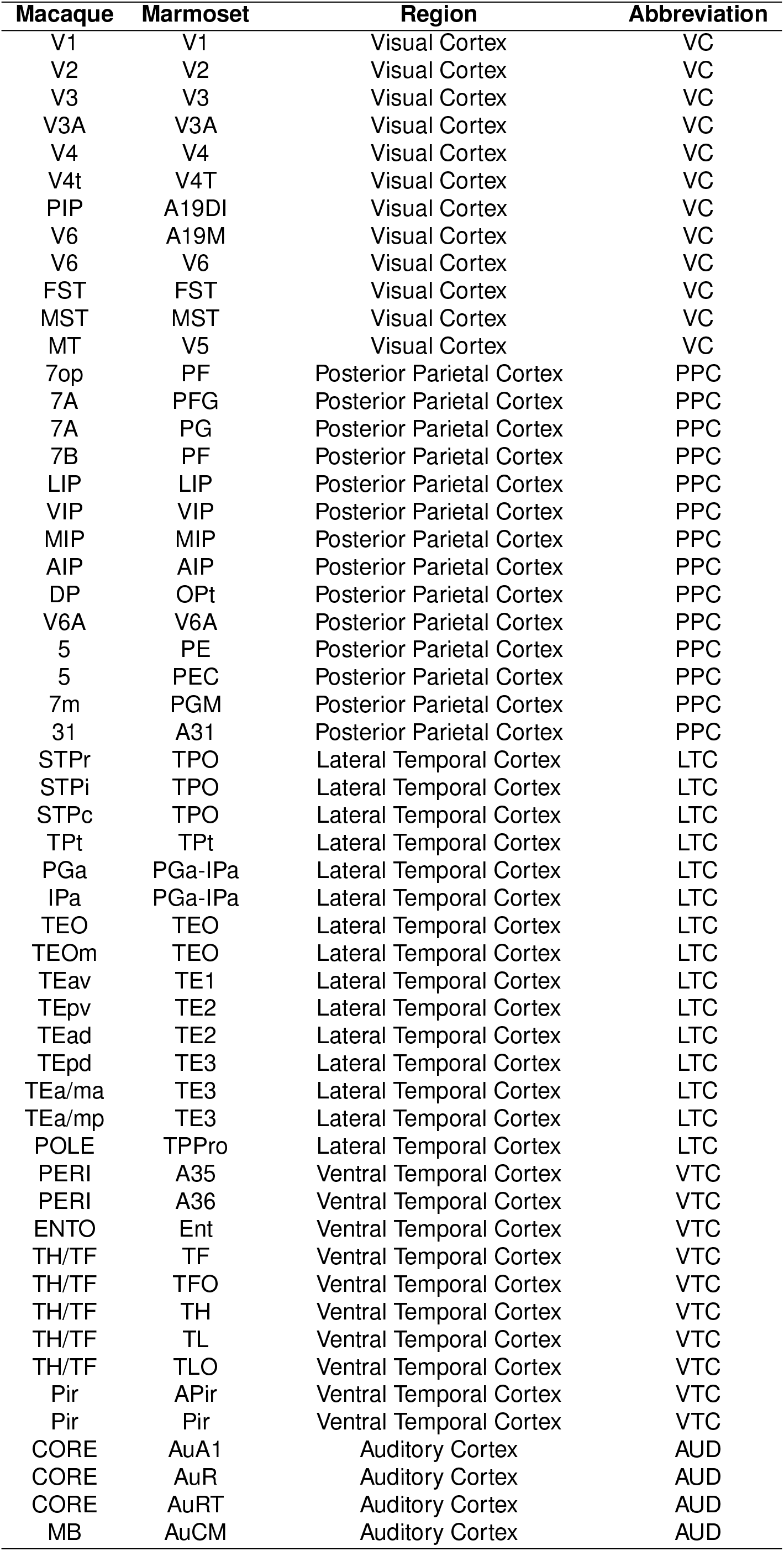
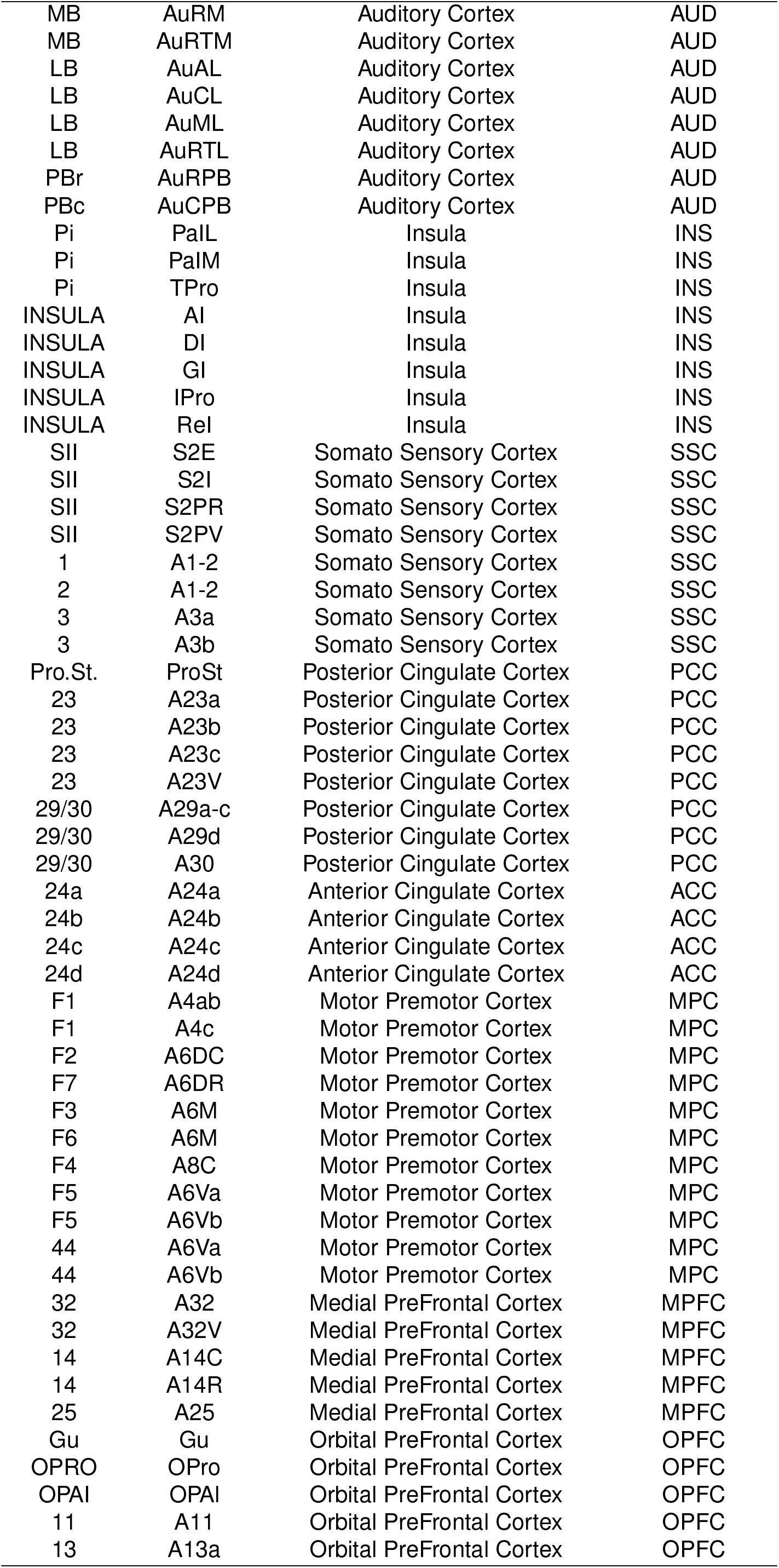
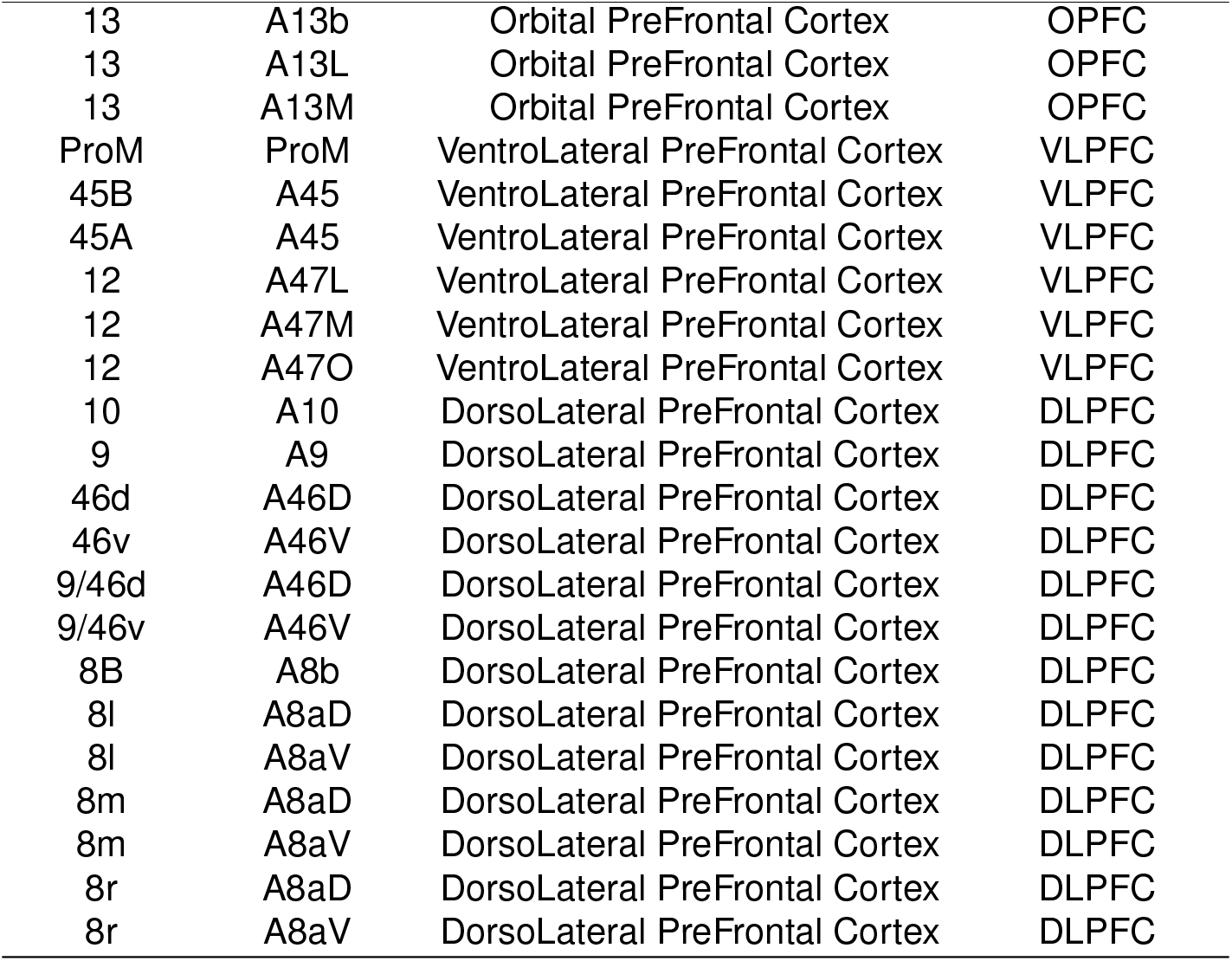
Consensus mapping atlas equivalence table. original atlases from Markov et al. ^1^ for the macaque and from Majka et al. ^2^ for the marmoset.

| **Macaque** | **Marmoset** | **Region** | **Abbreviation** |
| --- | --- | --- | --- |
| V1 | V1 | Visual Cortex | VC |
| V2 | V2 | Visual Cortex | VC |
| V3 | V3 | Visual Cortex | VC |
| V3A | V3A | Visual Cortex | VC |
| V4 | V4 | Visual Cortex | VC |
| V4t | V4T | Visual Cortex | VC |
| PIP | A19DI | Visual Cortex | VC |
| V6 | A19M | Visual Cortex | VC |
| V6 | V6 | Visual Cortex | VC |
| FST | FST | Visual Cortex | VC |
| MST | MST | Visual Cortex | VC |
| MT | V5 | Visual Cortex | VC |
| 7op | PF | Posterior Parietal Cortex | PPC |
| 7A | PFG | Posterior Parietal Cortex | PPC |
| 7A | PG | Posterior Parietal Cortex | PPC |
| 7B | PF | Posterior Parietal Cortex | PPC |
| LIP | LIP | Posterior Parietal Cortex | PPC |
| VIP | VIP | Posterior Parietal Cortex | PPC |
| MIP | MIP | Posterior Parietal Cortex | PPC |
| AIP | AIP | Posterior Parietal Cortex | PPC |
| DP | OPt | Posterior Parietal Cortex | PPC |
| V6A | V6A | Posterior Parietal Cortex | PPC |
| 5 | PE | Posterior Parietal Cortex | PPC |
| 5 | PEC | Posterior Parietal Cortex | PPC |
| 7m | PGM | Posterior Parietal Cortex | PPC |
| 31 | A31 | Posterior Parietal Cortex | PPC |
| STPr | TPO | Lateral Temporal Cortex | LTC |
| STPi | TPO | Lateral Temporal Cortex | LTC |
| STPc | TPO | Lateral Temporal Cortex | LTC |
| TPt | TPt | Lateral Temporal Cortex | LTC |
| PGa | PGa-IPa | Lateral Temporal Cortex | LTC |
| IPa | PGa-IPa | Lateral Temporal Cortex | LTC |
| TEO | TEO | Lateral Temporal Cortex | LTC |
| TEOm | TEO | Lateral Temporal Cortex | LTC |
| TEav | TE1 | Lateral Temporal Cortex | LTC |
| TEpv | TE2 | Lateral Temporal Cortex | LTC |
| TEad | TE2 | Lateral Temporal Cortex | LTC |
| TEpd | TE3 | Lateral Temporal Cortex | LTC |
| TEa/ma | TE3 | Lateral Temporal Cortex | LTC |
| TEa/mp | TE3 | Lateral Temporal Cortex | LTC |
| POLE | TPPro | Lateral Temporal Cortex | LTC |
| PERI | A35 | Ventral Temporal Cortex | VTC |
| PERI | A36 | Ventral Temporal Cortex | VTC |
| ENTO | Ent | Ventral Temporal Cortex | VTC |
| TH/TF | TF | Ventral Temporal Cortex | VTC |
| TH/TF | TFO | Ventral Temporal Cortex | VTC |
| TH/TF | TH | Ventral Temporal Cortex | VTC |
| TH/TF | TL | Ventral Temporal Cortex | VTC |
| TH/TF | TLO | Ventral Temporal Cortex | VTC |
| Pir | APir | Ventral Temporal Cortex | VTC |
| Pir | Pir | Ventral Temporal Cortex | VTC |
| CORE | AuA1 | Auditory Cortex | AUD |
| CORE | AuR | Auditory Cortex | AUD |
| CORE | AuRT | Auditory Cortex | AUD |
| MB | AuCM | Auditory Cortex | AUD |
| MB | AuRM | Auditory Cortex | AUD |
| MB | AuRTM | Auditory Cortex | AUD |
| LB | AuAL | Auditory Cortex | AUD |
| LB | AuCL | Auditory Cortex | AUD |
| LB | AuML | Auditory Cortex | AUD |
| LB | AuRTL | Auditory Cortex | AUD |
| PBr | AuRPB | Auditory Cortex | AUD |
| PBc | AuCPB | Auditory Cortex | AUD |
| Pi | PalL | Insula | INS |
| Pi | PalM | Insula | INS |
| Pi | TPro | Insula | INS |
| INSULA | AI | Insula | INS |
| INSULA | DI | Insula | INS |
| INSULA | GI | Insula | INS |
| INSULA | IPro | Insula | INS |
| INSULA | Rel | Insula | INS |
| SII | S2E | Somato Sensory Cortex | SSC |
| SII | S2I | Somato Sensory Cortex | SSC |
| SII | S2PR | Somato Sensory Cortex | SSC |
| SII | S2PV | Somato Sensory Cortex | SSC |
| 1 | A1-2 | Somato Sensory Cortex | SSC |
| 2 | A1-2 | Somato Sensory Cortex | SSC |
| 3 | A3a | Somato Sensory Cortex | SSC |
| 3 | A3b | Somato Sensory Cortex | SSC |
| Pro.St. | ProSt | Posterior Cingulate Cortex | PCC |
| 23 | A23a | Posterior Cingulate Cortex | PCC |
| 23 | A23b | Posterior Cingulate Cortex | PCC |
| 23 | A23c | Posterior Cingulate Cortex | PCC |
| 23 | A23V | Posterior Cingulate Cortex | PCC |
| 29/30 | A29a-c | Posterior Cingulate Cortex | PCC |
| 29/30 | A29d | Posterior Cingulate Cortex | PCC |
| 29/30 | A30 | Posterior Cingulate Cortex | PCC |
| 24a | A24a | Anterior Cingulate Cortex | ACC |
| 24b | A24b | Anterior Cingulate Cortex | ACC |
| 24c | A24c | Anterior Cingulate Cortex | ACC |
| 24d | A24d | Anterior Cingulate Cortex | ACC |
| F1 | A4ab | Motor Premotor Cortex | MPC |
| F1 | A4c | Motor Premotor Cortex | MPC |
| F2 | A6DC | Motor Premotor Cortex | MPC |
| F7 | A6DR | Motor Premotor Cortex | MPC |
| F3 | A6M | Motor Premotor Cortex | MPC |
| F6 | A6M | Motor Premotor Cortex | MPC |
| F4 | A8C | Motor Premotor Cortex | MPC |
| F5 | A6Va | Motor Premotor Cortex | MPC |
| F5 | A6Vb | Motor Premotor Cortex | MPC |
| 44 | A6Va | Motor Premotor Cortex | MPC |
| 44 | A6Vb | Motor Premotor Cortex | MPC |
| 32 | A32 | Medial PreFrontal Cortex | MPFC |
| 32 | A32V | Medial PreFrontal Cortex | MPFC |
| 14 | A14C | Medial PreFrontal Cortex | MPFC |
| 14 | A14R | Medial PreFrontal Cortex | MPFC |
| 25 | A25 | Medial PreFrontal Cortex | MPFC |
| Gu | Gu | Orbital PreFrontal Cortex | OPFC |
| OPRO | OPro | Orbital PreFrontal Cortex | OPFC |
| OPAI | OPAI | Orbital PreFrontal Cortex | OPFC |
| 11 | A11 | Orbital PreFrontal Cortex | OPFC |
| 13 | A13a | Orbital PreFrontal Cortex | OPFC |
| 13 | A13b | Orbital PreFrontal Cortex | OPFC |
| 13 | A13L | Orbital PreFrontal Cortex | OPFC |
| 13 | A13M | Orbital PreFrontal Cortex | OPFC |
| ProM | ProM | VentroLateral PreFrontal Cortex | VLPFC |
| 45B | A45 | VentroLateral PreFrontal Cortex | VLPFC |
| 45A | A45 | VentroLateral PreFrontal Cortex | VLPFC |
| 12 | A47L | VentroLateral PreFrontal Cortex | VLPFC |
| 12 | A47M | VentroLateral PreFrontal Cortex | VLPFC |
| 12 | A47O | VentroLateral PreFrontal Cortex | VLPFC |
| 10 | A10 | DorsoLateral PreFrontal Cortex | DLPFC |
| 9 | A9 | DorsoLateral PreFrontal Cortex | DLPFC |
| 46d | A46D | DorsoLateral PreFrontal Cortex | DLPFC |
| 46v | A46V | DorsoLateral PreFrontal Cortex | DLPFC |
| 9/46d | A46D | DorsoLateral PreFrontal Cortex | DLPFC |
| 9/46v | A46V | DorsoLateral PreFrontal Cortex | DLPFC |
| 8B | A8b | DorsoLateral PreFrontal Cortex | DLPFC |
| 8l | A8aD | DorsoLateral PreFrontal Cortex | DLPFC |
| 8l | A8aV | DorsoLateral PreFrontal Cortex | DLPFC |
| 8m | A8aD | DorsoLateral PreFrontal Cortex | DLPFC |
| 8m | A8aV | DorsoLateral PreFrontal Cortex | DLPFC |
| 8r | A8aD | DorsoLateral PreFrontal Cortex | DLPFC |
| 8r | A8aV | DorsoLateral PreFrontal Cortex | DLPFC |

**Table S4:** Marmoset pine count, hierarchical values and literary sources. Reading conventions as in **Table S1**. All parameter values are taken from Mejias and Wang ^3^ apart from *G* and *ρ*.

| Parameter | Value | Explanation |
| --- | --- | --- |
| **Current equation parameters** |  |  |
| $J_s$ | 0.3213 | Self-coupling current for E pop [nA] |
| $J_c$ | 0.0107 | Cross-coupling current for E pop [nA] |
| $J_{EI}$ | -0.31 | Coupling current from I to E pop [nA] |
| $J_{II}$ | -0.12 | Coupling current from I to I pop [nA] |
| $I_{0AB}$ | 0.3294 | Background current for E pop [nA] |
| $I_{0C}$ | 0.26 | Background current for I pop [nA] |
| **Conductance equation parameters** |  |  |
| $\tau_N$ | 0.06 | NMDA excitatory time constant [s] |
| $\tau_G$ | 0.005 | GABA inhibitory time constant [s] |
| $\gamma$ | 1.282 | Saturation factor for excitatory conductances [∅] |
| $\gamma_I$ | 2 | Saturation factor for inhibitory conductance [∅] |
| **Noise equation parameters** |  |  |
| $\tau_n$ | 0.002 | Noise time constant [s] |
| $\sigma_{AB}$ | 0.01 | Noise magnitude for excitatory pop A and B [nA] |
| $\sigma_C$ | 0 | Noise magnitude for inhibitory pop C [nA] |
| **Transfer function Parameters** |  |  |
| $a$ | 135 | Slope coefficient [Hz/nA] |
| $b$ | 54 | intercept [Hz] |
| $d$ | 0.308 | exponential coefficient [s] |
| $g_I$ | 4 | Descaling parameter [∅] |
| $c_0$ | 177 | Frequency threshold [Hz] |
| $c_1$ | 615 | Conversion slope [Hz/nA] |
| $r_0$ | 5.5 | intercept [Hz] |
| $\tau_r$ | 0.002 | Firing rate time constant [s] |
| **Synaptic strength parameters** |  |  |
| $J_{min}$ | 0.21 | Minimal synaptic strength [nA] |
| $J_{max}$ | 0.42 | Maximal synaptic strength [nA] |
| $J_0$ | 0.2112 | Basic synaptic strength for computing gradient of $J_{IE}$ [nA] |
| **Inter-areal connectivity** |  |  |
| $G$ | 0.98-0.85 | Global coupling strength [∅] |
| $k_1$ | 1.2 | FLN scaling parameter 1 [∅] |
| $k_2$ | 0.3 | FLN scaling parameter 2 [∅] |
| $\rho_1$ | 1 | FLN amplification from PFC to PFC [∅] |
| $\rho_2$ | 1 | FLN amplification from PFC to PPC [∅] |
| $\rho_3$ | 1 | FLN amplification from PPC to PPC [∅] |
| $\rho_4$ | 1 | FLN amplification from PPC to PFC [∅] |

### Downloadable supplementary material

Complete consensus meso-connectome data, original code and supplementary videos S1 and S2 has been deposited at Zenodo under the DOI 10.5281/zenodo.15612783

## References

1. Mayr, E. (1982). The Growth of Biological Thought: Diversity, Evolution, and Inheritance. Belknap Press.

2. Sporns, O., Tononi, G., and Kötter, R. (2005). The human connectome: a structural description of the human brain. PLoS computational biology 1, e42.

3. Hagmann, P. From diffusion mri to brain connectomics. Ph.D. thesis É cole Polytechnique Fédérale de Lausanne (EPFL) (2005).

4. Harris, K.D., and Mrsic-Flogel, T.D. (2013). Cortical connectivity and sensory coding. Nature 503, 51–58.

5. Binzegger, T., Douglas, R.J., and Martin, K.A. (2009). Topology and dynamics of the canonical circuit of cat v1. Neural Networks 22, 1071–1078.

6. Binzegger, T., Douglas, R.J., and Martin, K.A. (2004). A quantitative map of the circuit of cat primary visual cortex. Journal of Neuroscience 24, 8441–8453.

7. Douglas, R.J., Martin, K.A., and Whitteridge, D. (1989). A canonical microcircuit for neocortex. Neural computation 1, 480–488.

8. Theodoni, P., Majka, P., Reser, D.H., Wójcik, D.K., Rosa, M.G., and Wang, X.J. (2022). Structural attributes and principles of the neocortical connectome in the marmoset monkey. Cerebral Cortex 32, 15–28.

9. Majka, P., Bai, S., Bakola, S., Bednarek, S., Chan, J.M., Jermakow, N., Passarelli, L., Reser, D.H., Theodoni, P., Worthy, K.H. et al. (2020). Open access resource for cellular-resolution analyses of corticocortical connectivity in the marmoset monkey. Nature communications 11, 1133.

10. Dorkenwald, S., McKellar, C.E., Macrina, T., Kemnitz, N., Lee, K., Lu, R., Wu, J., Popovych, S., Mitchell, E., Nehoran, B. et al. (2022). Flywire: online community for whole-brain connectomics. Nature methods 19, 119–128.

11. Scheffer, L.K., Xu, C.S., Januszewski, M., Lu, Z., Takemura, S.y., Hayworth, K.J., Huang, G.B., Shinomiya, K., Maitlin-Shepard, J., Berg, S. et al. (2020). A connectome and analysis of the adult drosophila central brain. elife 9, e57443.

12. Harris, J.A., Mihalas, S., Hirokawa, K.E., Whitesell, J.D., Choi, H., Bernard, A., Bohn, P., Caldejon, S., Casal, L., Cho, A. et al. (2019). Hierarchical organization of cortical and thalamic connectivity. Nature 575, 195–202.

13. Gămănuţ, R., Kennedy, H., Toroczkai, Z., Ercsey-Ravasz, M., Van Essen, D.C., Knoblauch, K., and Burkhalter, A. (2018). The mouse cortical connectome, characterized by an ultradense cortical graph, maintains specificity by distinct connectivity profiles. Neuron 97, 698–715.

14. Markov, N.T., Ercsey-Ravasz, M.M., Ribeiro Gomes, A., Lamy, C., Magrou, L., Vezoli, J., Misery, P., Falchier, A., Quilodran, R., Gariel, M.A. et al. (2014). A weighted and directed interareal connectivity matrix for macaque cerebral cortex. Cerebral cortex 24, 17–36.

15. Markov, N.T., Vezoli, J., Chameau, P., Falchier, A., Quilodran, R., Huissoud, C., Lamy, C., Misery, P., Giroud, P., Ullman, S. et al. (2014). Anatomy of hierarchy: feedforward and feedback pathways in macaque visual cortex. Journal of comparative neurology 522, 225–259.

16. Felleman, D.J., and Van Essen, D.C. (1991). Distributed hierarchical processing in the primate cerebral cortex. Cerebral cortex (New York, NY: 1991) 1, 1–47.

17. Horvát, S., Gă mă nut, R., Ercsey-Ravasz, M., Magrou, L., Gă mă nut, B., Van Essen, D.C., Burkhalter, A., Knoblauch, K., Toroczkai, Z., and Kennedy, H. (2016). Spatial embedding and wiring cost constrain the functional layout of the cortical network of rodents and primates. PLoS biology 14, e1002512.

18. Ercsey-Ravasz, M., Markov, N.T., Lamy, C., Van Essen, D.C., Knoblauch, K., Toroczkai, Z., and Kennedy, H. (2013). A predictive network model of cerebral cortical connectivity based on a distance rule. Neuron 80, 184–197.

19. Magrou, L., Joyce, M.K.P., Froudist-Walsh, S., Datta, D., Wang, X.J., Martinez-Trujillo, J., and Arnsten, A.F. (2024). The meso-connectomes of mouse, marmoset, and macaque: network organization and the emergence of higher cognition. Cerebral Cortex 34, bhae174.

20. Wang, X.J., and Kennedy, H. (2016). Brain structure and dynamics across scales: in search of rules. Current opinion in neurobiology 37, 92–98.

21. Markov, N.T., Ercsey-Ravasz, M., Van Essen, D.C., Knoblauch, K., Toroczkai, Z., and Kennedy, H. (2013). Cortical high-density counterstream architectures. Science 342, 1238406.

22. Murre, J.M., and Sturdy, D.P. (1995). The connectivity of the brain: multi-level quantitative analysis. Biological cybernetics 73, 529–545.

23. Wang, X.J. (2022). Theory of the multiregional neocortex: large-scale neural dynamics and distributed cognition. Annual review of neuroscience 45, 533–560.

24. Sreenivasan, K.K., and D’Esposito, M. (2019). The what, where and how of delay activity. Nature reviews neuroscience 20, 466–481.

25. Christophel, T.B., Klink, P.C., Spitzer, B., Roelfsema, P.R., and Haynes, J.D. (2017). The distributed nature of working memory. Trends in cognitive sciences 21, 111–124.

26. Leavitt, M.L., Mendoza-Halliday, D., and Martinez-Trujillo, J.C. (2017). Sustained activity encoding working memories: not fully distributed. Trends in Neurosciences 40, 328–346.

27. Goldman-Rakic, P.S. (1992). Working memory and the mind. Scientific American 267, 110–117.

28. Fuster, J.M., and Alexander, G.E. (1971). Neuron activity related to short-term memory. Science 173, 652–654.

29. Wang, X.J. (2001). Synaptic reverberation underlying mnemonic persistent activity. Trends in neurosciences 24, 455–463.

30. Wang, X.J. (2021). 50 years of mnemonic persistent activity: quo vadis? Trends in Neurosciences 44, 888–902.

31. Mejias, J.F., and Wang, X.J. (2022). Mechanisms of distributed working memory in a largescale network of macaque neocortex. Elife 11, e72136.

32. Froudist-Walsh, S., Xu, T., Niu, M., Rapan, L., Zhao, L., Margulies, D.S., Zilles, K., Wang, X.J., and Palomero-Gallagher, N. (2023). Gradients of neurotransmitter receptor expression in the macaque cortex. Nature neuroscience 26, 1281–1294.

33. Chafee, M.V., and Goldman-Rakic, P.S. (1998). Matching patterns of activity in primate prefrontal area 8a and parietal area 7ip neurons during a spatial working memorytask. Journal of neurophysiology 79, 2919–2940.

34. Suzuki, M., and Gottlieb, J. (2013). Distinct neural mechanisms of distractor suppression in the frontal and parietal lobe. Nature neuroscience 16, 98–104.

35. Olesen, P.J., Macoveanu, J., Tegnér, J., and Klingberg, T. (2007). Brain activity related to working memory and distraction in children and adults. Cerebral cortex 17, 1047–1054.

36. Lorenc, E.S., Mallett, R., and Lewis-Peacock, J.A. (2021). Distraction in visual working memory: Resistance is not futile. Trends in cognitive sciences 25, 228–239.

37. Bisley, J.W., and Goldberg, M.E. (2006). Neural correlates of attention and distractibility in the lateral intraparietal area. Journal of neurophysiology 95, 1696–1717.

38. Lennert, T., and Martinez-Trujillo, J. (2011). Strength of response suppression to distracter stimuli determines attentional-filtering performance in primate prefrontal neurons. Neuron 70, 141–152.

39. Qi, X., Katsuki, F., Meyer, T., Rawley, J.B., Zhou, X., Douglas, K.L., and Constantinidis, C. (2010). Comparison of neural activity related to working memory in primate dorsolateral prefrontal and posterior parietal cortex. Frontiers in systems neuroscience 4, 1261.

40. Murray, J.D., Jaramillo, J., and Wang, X.J. (2017). Working memory and decision-making in a frontoparietal circuit model. Journal of Neuroscience 37, 12167–12186.

41. Prins, N.W., Pohlmeyer, E.A., Debnath, S., Mylavarapu, R., Geng, S., Sanchez, J.C., Rothen, D., and Prasad, A. (2017). Common marmoset (callithrix jacchus) as a primate model for behavioral neuroscience studies. Journal of neuroscience methods 284, 35–46.

42. Mitchell, J.F., and Leopold, D.A. (2015). The marmoset monkey as a model for visual neuroscience. Neuroscience research 93, 20–46.

43. Okano, H., Hikishima, K., Iriki, A., and Sasaki, E. (2012). The common marmoset as a novel animal model system for biomedical and neuroscience research applications. In Seminars in fetal and neonatal medicine vol. 17. Elsevier pp. 336–340.

44. Reser, D.H., Burman, K.J., Yu, H.H., Chaplin, T.A., Richardson, K.E., Worthy, K.H., and Rosa, M.G. (2013). Contrasting patterns of cortical input to architectural subdivisions of the area 8 complex: a retrograde tracing study in marmoset monkeys. Cerebral Cortex 23, 1901–1922.

45. Spinelli, S., Pennanen, L., Dettling, A.C., Feldon, J., Higgins, G.A., and Pryce, C.R. (2004). Performance of the marmoset monkey on computerized tasks of attention and working memory. Cognitive Brain Research 19, 123–137.

46. Wong, R.K., Selvanayagam, J., Johnston, K.D., and Everling, S. (2023). Delay-related activity in marmoset prefrontal cortex. Cerebral Cortex 33, 3523–3537.

47. Nakamura, K., Koba, R., Miwa, M., Yamaguchi, C., Suzuki, H., and Takemoto, A. (2018). A method to train marmosets in visual working memory task and their performance. Frontiers in behavioral neuroscience 12, 46.

48. Zlatkina, V., Frey, S., and Petrides, M. (2024). Monitoring of nonspatial information within working memory in the common marmoset (callithrix jacchus). Cerebral Cortex 34, bhae444.

49. Joyce, M.K.P., Ivanov, T.G., Krienen, F.M., Mitchell, J.F., Ma, S., Inoue, W., Nandy, A.S., Datta, D., Duque, A., Arellano, J.I. et al. (2025). Higher dopamine d1 receptor expression in prefrontal parvalbumin neurons underlies higher distractibility in marmosets versus macaques. Communications Biology 8, 974.

50. Markov, N.T., Misery, P., Falchier, A., Lamy, C., Vezoli, J., Quilodran, R., Gariel, M., Giroud, P., Ercsey-Ravasz, M., Pilaz, L. et al. (2011). Weight consistency specifies regularities of macaque cortical networks. Cerebral cortex 21, 1254–1272.

51. Vezoli, J., Magrou, L., Goebel, R., Wang, X.J., Knoblauch, K., Vinck, M., and Kennedy, H. (2021). Cortical hierarchy, dual counterstream architecture and the importance of topdown generative networks. Neuroimage 225, 117479.

52. Delhaye, B.P., Long, K.H., and Bensmaia, S.J. (2018). Neural basis of touch and proprioception in primate cortex. Comprehensive Physiology 8, 1575.

53. Chaudhuri, R., Knoblauch, K., Gariel, M.A., Kennedy, H., and Wang, X.J. (2015). A largescale circuit mechanism for hierarchical dynamical processing in the primate cortex. Neuron 88, 419–431.

54. Bisley, J.W., and Goldberg, M.E. (2003). Neuronal activity in the lateral intraparietal area and spatial attention. Science 299, 81–86.

55. Jacob, S.N., and Nieder, A. (2014). Complementary roles for primate frontal and parietal cortex in guarding working memory from distractor stimuli. Neuron 83, 226–237.

56. Moeller, S., Nallasamy, N., Tsao, D.Y., and Freiwald, W.A. (2009). Functional connectivity of the macaque brain across stimulus and arousal states. Journal of Neuroscience 29, 5897–5909.

57. Flavell, S.W., Gogolla, N., Lovett-Barron, M., and Zelikowsky, M. (2022). The emergence and influence of internal states. Neuron 110, 2545–2570.

58. Greene, A.S., Horien, C., Barson, D., Scheinost, D., and Constable, R.T. (2023). Why is everyone talking about brain state? Trends in Neurosciences 46, 508–524.

59. Kringelbach, M.L., and Deco, G. (2020). Brain states and transitions: insights from computational neuroscience. Cell Reports 32.

60. Froudist-Walsh, S., Bliss, D.P., Ding, X., Rapan, L., Niu, M., Knoblauch, K., Zilles, K., Kennedy, H., Palomero-Gallagher, N., and Wang, X.J. (2021). A dopamine gradient controls access to distributed working memory in the large-scale monkey cortex. Neuron 109, 3500–3520.

61. Ringo, J.L. (1991). Neuronal interconnection as a function of brain size. Brain, behavior and evolution 38, 1–6.

62. Changeux, J.P., Goulas, A., and Hilgetag, C.C. (2021). A connectomic hypothesis for the hominization of the brain. Cerebral cortex 31, 2425–2449.

63. Du, J., DiNicola, L.M., Angeli, P.A., Saadon-Grosman, N., Sun, W., Kaiser, S., Ladopoulou, J., Xue, A., Yeo, B.T., Eldaief, M.C. et al. (2024). Organization of the human cerebral cortex estimated within individuals: networks, global topography, and function. Journal of Neurophysiology 131, 1014–1082.

64. Blanchard, P., Devaney, R., and Hall, G. (2012). Differential Equations. Cengage Learning. ISBN 9780495561989.

65. Balasubramaniam, K.N., Dittmar, K., Berman, C.M., Butovskaya, M., Cooper, M.A., Majolo, B., Ogawa, H., Schino, G., Thierry, B., and De Waal, F.B. (2012). Hierarchical steepness, counter-aggression, and macaque social style scale. American journal of primatology 74, 915–925.

66. Sehner, S., Willems, E.P., Vinicus, L., Migliano, A.B., van Schaik, C.P., and Burkart, J.M. (2022). Problem-solving in groups of common marmosets (callithrix jacchus): more than the sum of its parts. PNAS nexus 1, pgac168.

67. Miller, C.T., Freiwald, W.A., Leopold, D.A., Mitchell, J.F., Silva, A.C., and Wang, X. (2016). Marmosets: a neuroscientific model of human social behavior. Neuron 90, 219–233.

68. Wong, K.F., and Wang, X.J. (2006). A recurrent network mechanism of time integration in perceptual decisions. Journal of Neuroscience 26, 1314–1328.

69. Gajwani, M., Oldham, S., Pang, J.C., Arnatkevičiūtė, A., Tiego, J., Bellgrove, M.A., and Fornito, A. (2023). Can hubs of the human connectome be identified consistently with diffusion mri? Network Neuroscience 7, 1326–1350.

70. Consagra, W., Venkataraman, A., and Zhang, Z. (2022). Optimized diffusion imaging for brain structural connectome analysis. IEEE transactions on medical imaging 41, 2118– 2129.

71. Donahue, C.J., Sotiropoulos, S.N., Jbabdi, S., Hernandez-Fernandez, M., Behrens, T.E., Dyrby, T.B., Coalson, T., Kennedy, H., Knoblauch, K., Van Essen, D.C. et al. (2016). Using diffusion tractography to predict cortical connection strength and distance: a quantitative comparison with tracers in the monkey. Journal of Neuroscience 36, 6758–6770.

72. Palmer, S.M., and Rosa, M.G. (2006). Quantitative analysis of the corticocortical projections to the middle temporal area in the marmoset monkey: evolutionary and functional implications. Cerebral Cortex 16, 1361–1375.

73. Burman, K.J., Bakola, S., Richardson, K.E., Reser, D.H., and Rosa, M.G. (2014). Patterns of cortical input to the primary motor area in the marmoset monkey. Journal of Comparative Neurology 522, 811–843.

74. Burman, K.J., Bakola, S., Richardson, K.E., Reser, D.H., and Rosa, M.G. (2014). Patterns of afferent input to the caudal and rostral areas of the dorsal premotor cortex (6dc and 6dr) in the marmoset monkey. Journal of Comparative Neurology 522, 3683–3716.

75. Falchier, A., Clavagnier, S., Barone, P., and Kennedy, H. (2002). Anatomical evidence of multimodal integration in primate striate cortex. Journal of Neuroscience 22, 5749–5759.

76. Barone, P., Batardiere, A., Knoblauch, K., and Kennedy, H. (2000). Laminar distribution of neurons in extrastriate areas projecting to visual areas v1 and v4 correlates with the hierarchical rank and indicates the operation of a distance rule. Journal of Neuroscience 20, 3263–3281.

77. Elston, G.N., Benavides-Piccione, R., Elston, A., Manger, P.R., and DeFelipe, J. (2011). Pyramidal cells in prefrontal cortex of primates: marked differences in neuronal structure among species. Front. Neuroanat. 5, 2.

78. Elston, G.N., Oga, T., Okamoto, T., and Fujita, I. (2010). Spinogenesis and pruning from early visual onset to adulthood: an intracellular injection study of layer iii pyramidal cells in the ventral visual cortical pathway of the macaque monkey. Cerebral Cortex 20, 1398– 1408.

79. Elston, G.N., Oga, T., and Fujita, I. (2009). Spinogenesis and pruning scales across functional hierarchies. Journal of Neuroscience 29, 3271–3275.

80. Elston, G.N., and Rockland, K.S. (2002). The pyramidal cell of the sensorimotor cortex of the macaque monkey: phenotypic variation. Cerebral Cortex 12, 1071–1078.

81. Elston, G. (2001). Interlaminar differences in the pyramidal cell phenotype in cortical areas 7m and stp (the superior temporal polysensory area) of the macaque monkey. Experimental brain research 138, 141–152.

82. Elston, G.N., Benavides-Piccione, R., and DeFelipe, J. (2001). The pyramidal cell in cognition: a comparative study in human and monkey. The Journal of Neuroscience 21, RC163.

83. Elston, G.N., and Rosa, M.G. (1998). Complex dendritic fields of pyramidal cells in the frontal eye field of the macaque monkey: comparison with parietal areas 7a and lip. Neuroreport 9, 127–131.

84. Elston, G.N., Tweedale, R., and Rosa, M.G.P. (1999). Cortical integration in the visual system of the macaque monkey: large-scale morphological differences in the pyramidal neurons in the occipital, parietal and temporal lobes. Proceedings of the Royal Society of London. Series B: Biological Sciences 266, 1367–1374.

85. Elston, G.N., and Rosa, M. (1998). Morphological variation of layer iii pyramidal neurones in the occipitotemporal pathway of the macaque monkey visual cortex. Cerebral cortex (New York, NY: 1991) 8, 278–294.

86. Elston, G.N., and Rosa, M. (1997). The occipitoparietal pathway of the macaque monkey: comparison of pyramidal cell morphology in layer iii of functionally related cortical visual areas. Cerebral cortex (New York, NY: 1991) 7, 432–452.

87. Oga, T., Aoi, H., Sasaki, T., Fujita, I., and Ichinohe, N. (2013). Postnatal development of layer iii pyramidal cells in the primary visual, inferior temporal, and prefrontal cortices of the marmoset. Frontiers in neural circuits 7, 31.

88. Sasaki, T., Aoi, H., Oga, T., Fujita, I., and Ichinohe, N. (2015). Postnatal development of dendritic structure of layer iii pyramidal neurons in the medial prefrontal cortex of marmoset. Brain Structure and Function 220, 3245–3258.

89. Elston, G.N., Tweedale, R., and Rosa, M.G. (1999). Cellular heterogeneity in cerebral cortex: a study of the morphology of pyramidal neurones in visual areas of the marmoset monkey. Journal of Comparative Neurology 415, 33–51.

90. Nimchinsky, E.A., Sabatini, B.L., and Svoboda, K. (2002). Structure and function of dendritic spines. Annual review of physiology 64, 313–353.

91. Elston, G.N. (2007). Specialization of the neocortical pyramidal cell during primate evolution. Evolution of nervous systems pp. 191–242.

92. Van Essen, D.C., and Maunsell, J.H. (1983). Hierarchical organization and functional streams in the visual cortex. Trends in neurosciences 6, 370–375.

93. Rockland, K.S., and Pandya, D.N. (1979). Laminar origins and terminations of cortical connections of the occipital lobe in the rhesus monkey. Brain research 179, 3–20.

94. Paxinos, G., Watson, C., Petrides, M., Rosa, M., and Tokuno, H. (2012). The marmoset brain in stereotaxic coordinates. Elsevier Academic Press.

95. Solomon, S.G., and Rosa, M.G. (2014). A simpler primate brain: the visual system of the marmoset monkey. Frontiers in neural circuits 8, 96.

96. Bakola, S., Burman, K.J., and Rosa, M.G. (2015). The cortical motor system of the marmoset monkey (callithrix jacchus). Neuroscience research 93, 72–81.

97. Abbott, L.F., and Chance, F.S. (2005). Drivers and modulators from push-pull and balanced synaptic input. Progress in brain research 149, 147–155.

98. Tsushima, Y., Sasaki, Y., and Watanabe, T. (2006). Greater disruption due to failure of inhibitory control on an ambiguous distractor. Science 314, 1786–1788.

99. Hupé, J., James, A., Payne, B., Lomber, S., Girard, P., and Bullier, J. (1998). Cortical feedback improves discrimination between figure and background by v1, v2 and v3 neurons. Nature 394, 784–787.

100. Girard, P., and Bullier, J. (1989). Visual activity in area v2 during reversible inactivation of area 17 in the macaque monkey. Journal of neurophysiology 62, 1287–1302.

101. Lesnoff, M., and Lancelot, R. aods3: Analysis of Overdispersed Data using S3 Methods (2024). URL: https://CRAN.R-project.org/package=aods3 r package version 0.5.

## References

1. Markov, N.T., Ercsey-Ravasz, M.M., Ribeiro Gomes, A., Lamy, C., Magrou, L., Vezoli, J., Misery, P., Falchier, A., Quilodran, R., Gariel, M.A. et al. (2014). A weighted and directed interareal connectivity matrix for macaque cerebral cortex. Cerebral cortex 24, 17–36.

2. Majka, P., Bai, S., Bakola, S., Bednarek, S., Chan, J.M., Jermakow, N., Passarelli, L., Reser, D.H., Theodoni, P., Worthy, K.H. et al. (2020). Open access resource for cellular-resolution analyses of corticocortical connectivity in the marmoset monkey. Nature communications 11, 1133.

3. Mejias, J.F., and Wang, X.J. (2022). Mechanisms of distributed working memory in a largescale network of macaque neocortex. Elife 11, e72136.

